# Machine-guided design of synthetic cell type-specific *cis*-regulatory elements

**DOI:** 10.1101/2023.08.08.552077

**Authors:** SJ Gosai, RI Castro, N Fuentes, JC Butts, S Kales, RR Noche, K Mouri, PC Sabeti, SK Reilly, R Tewhey

## Abstract

*Cis*-regulatory elements (CREs) control gene expression, orchestrating tissue identity, developmental timing, and stimulus responses, which collectively define the thousands of unique cell types in the body. While there is great potential for strategically incorporating CREs in therapeutic or biotechnology applications that require tissue specificity, there is no guarantee that an optimal CRE for an intended purpose has arisen naturally through evolution. Here, we present a platform to engineer and validate synthetic CREs capable of driving gene expression with programmed cell type specificity. We leverage innovations in deep neural network modeling of CRE activity across three cell types, efficient *in silico* optimization, and massively parallel reporter assays (MPRAs) to design and empirically test thousands of CREs. Through *in vitro* and *in vivo* validation, we show that synthetic sequences outperform natural sequences from the human genome in driving cell type-specific expression. Synthetic sequences leverage unique sequence syntax to promote activity in the on-target cell type and simultaneously reduce activity in off-target cells. Together, we provide a generalizable framework to prospectively engineer CREs and demonstrate the required literacy to write regulatory code that is fit-for-purpose *in vivo* across vertebrates.

## Introduction

Our understanding of how CREs impact gene expression has been primarily derived from those elements that exist naturally in the human genome^1–4^. Major efforts over the past decade have identified millions of putative CREs, yet these sequences generated by evolution represent only a small subset of possible genetic sequences and may not meet expression objectives favorable for therapeutic applications^5–7^. Indeed, 200 base pairs of DNA can encompass over 2.58×10^120^ possible sequences, more combinations than atoms in the observable universe. This unexplored CRE sequence space, combined with our current poor understanding of the underlying principles driving CRE function, limit our ability to leverage CREs for clinical or biotechnological applications^8,9^. Bridging the gap in knowledge of ‘regulatory grammar’—the syntax of activating and repressing transcription factor (TF) vocabularies, their combinatorial effects, and higher order rules of TF cooperativity—has been a major goal of genomics for the past decade^6,7,10–13^.

Recent advances are reshaping our ability to design CRE sequences with cell type-specific activity by overcoming three gaps in knowledge: (1) scalable methods to functionally characterize natural and synthetic CREs to produce generalizable insights (2) accurate ‘regulatory grammar’ models of how genetic sequences lead to CRE activity across cell types, and (3) the ability to repurpose predictive models for directed CRE generation. First, MPRAs can directly characterize CRE activity potential at-scale and across cell types^14–19^. Hundreds of thousands of CREs have been functionally characterized by MPRA, providing initial insights into regulatory syntax and transcriptional specificity^20–24^. Second, deep learning has emerged as an effective tool to accurately model the relationship between genetic sequences and biological phenotypes^25–33^. While these sequence models are promising tools for the interpretation of genetic sequences^28,29,32,34^, they have largely been trained on, and predict, proxies of regulatory activity such as regions of open chromatin demarcated by DNAse Hypersensitivity sites (DHS), rather than direct CRE activity. Lastly, although computational models are millions of times faster than experimentation, they are still only capable of characterizing a fraction of all possible sequence combinations. Efficient frameworks to generate sequences from predictive models could enable rational and interpretable design of candidate CREs^9,35–40^.

Programmed, highly precise, cell type-specific transcriptional control CREs would contribute to development of specialized reporters, CRISPR therapeutics, gene replacement approaches, and more. In particular, advances in gene therapies offer a route to ameliorating a rapidly growing list of human genetic diseases, but their widespread use is hindered by a lack of robust, cell type-targeted delivery^41^. While current nanoparticle^42^ and viral vector^43^ technologies have shown some promise in better targeting of clinically actionable tissues like brain and muscle, they often display many undesirable cell type off-target effects^44,45^. Being able to fabricate synthetic CREs with programmable, highly tissue-specific functions could provide orthogonal tools for such clinical applications as well as basic research.

Here we present a method to engineer novel synthetic CREs capable of driving gene expression with cell type specificity. We leverage innovations in modeling regulatory grammar across cell types, efficient sequence space searching, and the MPRA experimental system that can validate thousands of CREs in parallel. We use a recently generated database of uniformly processed MPRA experiments which characterized an unprecedented number of CREs to train an accurate deep-learning model that can rapidly predict activity for any sequence *in silico*.

Coupled to sequence generation algorithms, we deploy our model to generate thousands of cell type-specific, synthetic CREs, which we functionally validate using MPRAs and *in vivo* using mouse and zebrafish.

## Results

### Deep learning models can accurately predict DNA *cis*-regulatory activity

We first built an accurate model of CRE activity from DNA sequence alone (**Figure 1a**). While previous models of CRE activity have primarily used epigenetic states correlated to CRE function^29,30,34,46,47^, we trained our model on the regulatory output of 776,474 200-nucleotide sequences directly, as assayed by MPRA, a high-throughput reporter system that quantifies the effect of a given sequence on gene transcription (**Supplementary Tables 1 and 2, Methods**). These MPRAs were conducted by a single lab using a consistent experimental and analytical pipeline, yielding highly reproducible measurements (**Supplementary Figure 1, Supplementary Table 2** ^24^, **Figure 1b**). In total, we collected functional CRE measurements from 155.3 Mbp of unique genomic sequence in each of three human cell types: K562 (erythroid precursors), HepG2 (hepatocytes), and SK-N-SH (neuroblastoma). These well-studied cell types are ideal for high-throughput method development and can provide useful insight for the growing body of experimental gene therapies that target blood cells^48–51^ and neurons^52^, but that can induce toxicity in the liver^53–55^.

**Figure 1.**
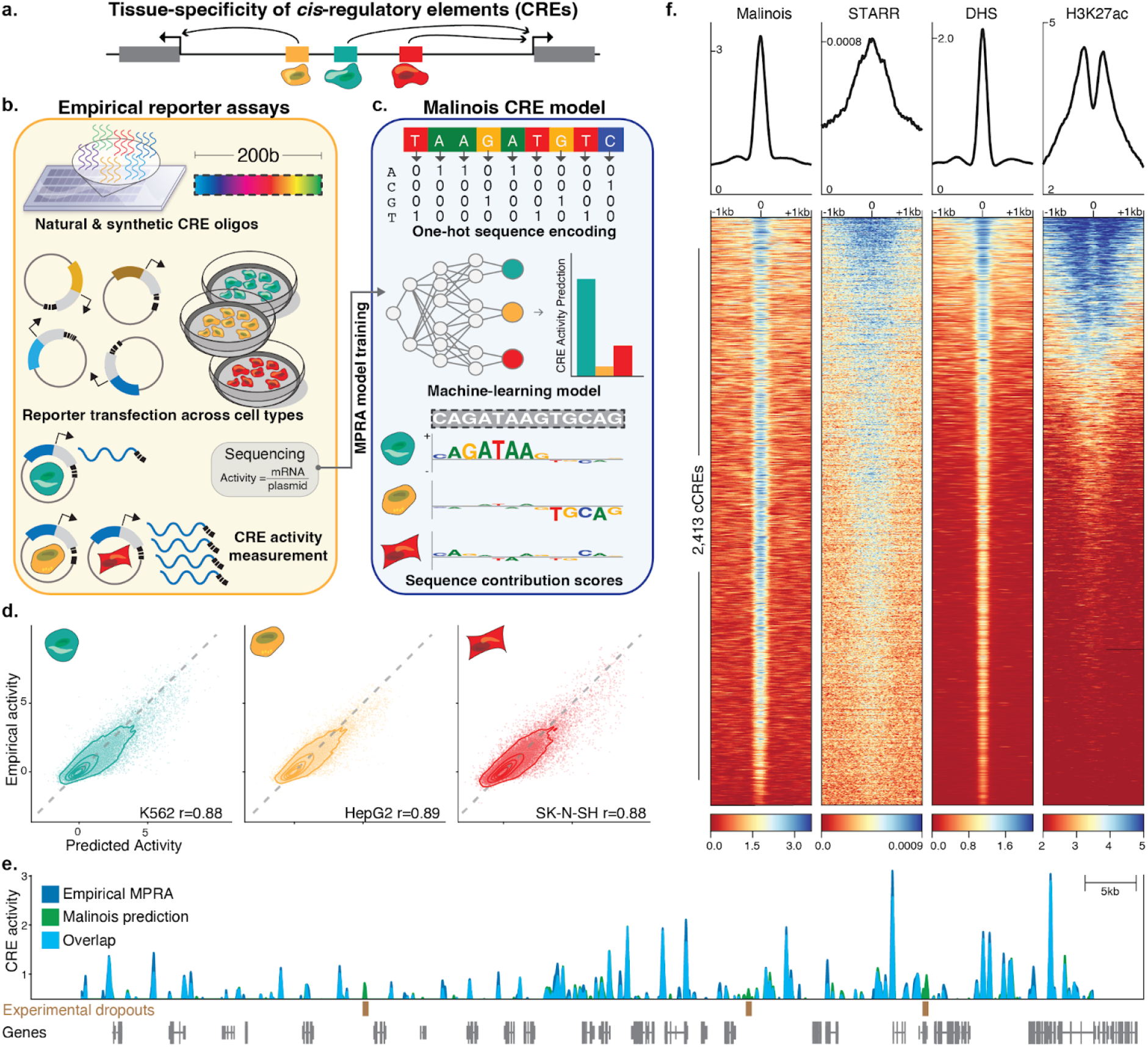
Malinois accurately predicts transcriptional activation by CREs in episomal reporters. (**a**) Schematic showing non-coding *cis*-regulatory elements (CREs) in the genome drive gene expression and contribute to cell type specific expression. (**b**) Overview of how MPRAs enable targeted functional characterization of hundreds of thousands of CREs on transcription in episomal reporters, and can quantify the impact of programmable 200-bp oligonucleotide sequences. MPRAs across multiple cell types enables discovery of cell type-specific activity of CREs. (**c**) Schematic showing how deep learning enables modeling of cell type-specific CRE effects directly from nucleotide sequence. Malinois, a deep convolutional neural network, predicts CRE activity in K562 (teal), HepG2 (yellow), and SK-N-SH (red). Contribution scores can be extracted from the model to determine how subsequences drive predicted function in each cell type. (**d**) Malinois predictions are highly correlated with empirically measured MPRA activity across K562 (teal), HepG2 (yellow), and SK-N-SH (red). Performance for each cell type was measured using Pearson correlation (r) on a test set of sequences withheld from training. Each point corresponds to empirical and predicted activity of a single CRE in the corresponding cell type, and topological lines indicate point density (16.7%, 33.3%, 50%, 66.7%, 83.3%) in the scatter plots. Train/test splits were defined by chromosomes. (**e**) Malinois activity predictions for sequences centered on K562-specific DHS peaks activate transcription in K562. This pattern of activation is concordant with quantitative signals measured using STARR-seq, DHS-seq, and H3K27ac seq. (**f**) Malinois predictions recapitulate an MPRA screen of overlapping fragments derived from a 2.1Mb window centered on the GATA1 gene (Pearson’s *r* = 0.91; **Supplementary Fig. 4**). Light blue signal indicates overlapping signal while dark blue and green regions indicate either higher activity measurements or predictions by MPRA or Malinois, respectively, in the window chrX:48,000,000-49,000,000.

We created Malinois, a deep convolutional neural network (CNN) for cell type-informed CRE activity prediction of any arbitrary sequence (**Figure 1c, Supplementary Figure 2, Methods**). We leveraged Bayesian optimization^56,57^ to iterate over one thousand CNN configurations and hyperparameter settings to identify the best model. Malinois accurately models episomal CRE activity across cell types. For sequences held out from training (62,582 elements on chromosomes 7 and 13), Malinois predictions in K562, HepG2, and SK-N-SH correlate highly with empirical activity measurements (Pearson’s *r* 0.88-0.89; Spearman’s ⍴ 0.81-0.83) (**Figure 1d**) and demonstrate cell specificity on par with experimental results (**Supplementary Figure 3**).

Given Malinois can accurately and rapidly model CRE activity, we generated genome-wide predictions of sequence activity to compare with orthogonal approaches for characterizing CREs. We observe a strong correlation (Pearson’s *r* = 0.91) between Malinois predictions and a comprehensive MPRA of sequences tiling a 2.1Mb window encompassing *GATA1* (**Figure 1e, Supplementary Figure 4**). We also find Malinois K562 predictions have strong activity at known markers of CREs identified by DHS sites^58^ (*p*<10^-300^, two-sided paired *t*-test) and H3K27ac ChIP-seq peaks^59,60^ (*p*<10^-114^, two-sided paired *t*-test), and are correlated with STARR-seq peaks^59,61^ (*p*<10^-178^, two-sided paired *t*-test), an orthogonal measure of CRE activity (**Figure 1f, Supplementary Figure 5, Supplementary Table 1**)^5,62–64^. This finding is consistent in HepG2 and SK-S-SH cells as well (**Supplementary Figure 5**). Together, this suggests Malinois predictions provide accurate measurements of CREs, approaching the biological reproducibility of empirical measures.

### CODA designs CREs with desired functions

We next developed CODA (Computational Optimization of DNA Activity), a modular platform for designing novel CREs with programmed functionality. CODA follows an iterative loop of predicting the activity of sequences, calculating a fitness value that quantifies how well sequences fit the design goals, and then updating sequences to improve fitness. Here, our design goal is cell type-specific CRE activity. Sequence updates in CODA can be controlled using different classes of sequence design algorithms. We implemented three classes of algorithms (evolutionary: AdaLead^37^, probabilistic: Simulated Annealing^65^, and gradient-based: Fast SeqProp^36^) for sequence generation. We selected these methodologies based on their ease of implementation, useful optimization guarantees, or their ability to exploit the structure of deep-learning models. Here, CODA uses Malinois as a fast and accurate measure of CRE activity, providing a scalable model to test billions of CRE designs within the optimization loop.

We deployed CODA to rationally design CREs with cell type-specific activity in K562, HepG2, and SK-N-SH cell lines (**Figure 2a**). This process involves six steps. We: (i) generate a set of random 200-mer sequences; (ii) predict regulatory activity of each sequence, in each cell type, using Malinois; (iii) transform these predictions using an objective function into a single fitness value of cell specificity; (iv) traverse the fitness landscape towards specificity by (v) modifying the sequence set *in silico* using one of the design algorithms (**Supplementary Figure 6**); and (vi) continue iterating until a batch of designed sequences reaches a fitness plateau. We define fitness as a function of the gap observed between predicted activity in the targeted cell type and the maximum of the two off-target cell types, herein referred to as MinGap (**Methods**).

**Figure 2.**
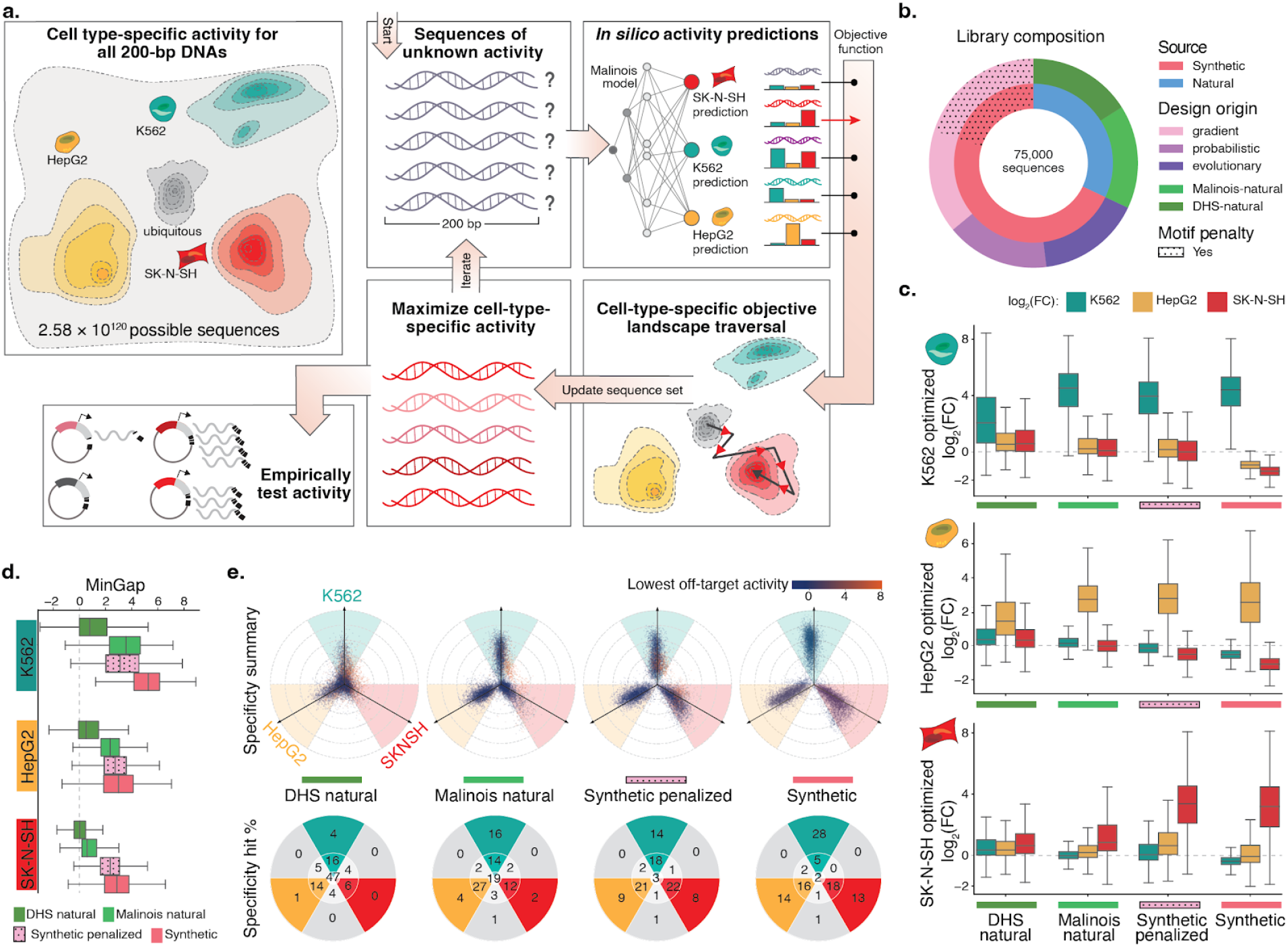
CODA effectively designs novel cell type-specific CREs using Malinois predictions. (**a**) CODA designs synthetic elements by iteratively updating sequences to improve predicted function. Cell type-specific CRE activity of all 200 bp DNA oligos induces a topology over a massive sample space. CODA initializes sequences in this space and uses Malinois to predict local topology. An objective function is used by CODA to direct updates of sequences to move as desired through predicted topology. Updated sequences can be further modified *in silico* until a stopping criteria is reached and final candidates are proposed for experimental validation. (**b**) Composition of the MPRA library designed to empirically evaluate candidate cell type-specific CREs. A total of 75,000 sequences were selected from the human genome (green hues) or designed *ab initio* using CODA (purple hues) to maximize the MinGap score for a target cell type. Aggregated natural and synthetic sequences are indicated by blue and coral coloring, respectively. Sequences generated using motif-penalization are delineated by the dotted overlay. (**c**) Computationally-designed CREs maintain high transcriptional activity in target cells while improving silencing in off-target cells. The three rows of box plots correspond to candidate CREs intended to drive cell type-specific expression in K562, HepG2, and SK-N-SH. Each group of three boxes indicate the distribution of MPRA log_2_ fold change (log_2_FC) measurements in K562 (teal), HepG2 (yellow), and SK-N-SH (red) for a set of sequences nominated by the indicated design strategy on the *x*-axis. Boxes demarcate the 25th, 50th, and 75th percentile values, while whiskers indicate the outermost point with 1.5 times the interquartile range from the edges of the boxes. Sequences with a replicate log_2_FC standard error greater than 1 in any cell type were not included. (**d**) CODA-designed synthetic sequences achieve higher overall cell type-specific activity than natural sequences. Box plots display distribution of MinGap scores to quantify cell-specific CRE function and color indicates intended target cell type (K562: teal; HepG2: yellow; SK-N-SH: red). Boxes demarcate the 25th, 50th, and 75th percentile values, while whiskers indicate the outermost point with 1.5 times the interquartile range from the edges of the boxes. Sequences with a replicate log_2_FC standard error greater than 1 in any cell type were not included. (**e**) Top row: propeller plots for each sequence group. The radial distance corresponds to the distance between the maximum and minimum cell type activity values, while the angle of deviation from an axis quantifies the relative activity of the highest off-target cell type (**Methods**). Teal, yellow, and red areas represent sequences in which the MinGap:MaxGap ratio is greater than 0.5. Dot colors are associated with the activity in the minimum off-target cell type. Bottom row: percentages of points in each delimited area rounded to the nearest integer. The point count in the center represents sequences with quasi-uniform activity across cell types, while the gray wedges count sequences with a low MinGap. The groups synthetic and synthetic-penalized were randomly sub-sampled to match the size of the two natural groups (see **Supplementary Fig 13** for full plots).

To empirically test the effectiveness of CODA, we performed an MPRA to measure activity of the synthetic sequences. For each cell type, we generated 4,000 cell type-specific sequences from each of the three sequence design algorithms in CODA, yielding a total of 36,000 synthetic candidates (**Figure 2b, Supplementary Table 3, Methods**). We observed that Malinois induced strong preferences for certain sequence motifs when maximizing specificity (**Supplementary Tables 4 and 5, Supplementary Figure 7a**). For this reason, we decided to also explore alternative solutions by encouraging CODA to modify the utilization of highly preferred motifs despite the potential decrease in predicted cell type specificity (**Methods**). Using Fast SeqProp, we designed a second group of synthetic sequences with a motif penalty incorporated into the fitness function (**Figure 2b**). Over five iterative rounds, we generated a total of 15,000 ’synthetic-penalized’ CREs, with 1,000 sequences per round per cell type, while penalizing the top motifs from the preceding rounds in each iteration (**Supplementary Table 4**). We observed successful reduction in initially enriched motifs and a simultaneous increase in motifs underutilized in earlier rounds (**Supplementary Figure 7b**), diversifying the syntax of CODA-proposed sequences for experimental evaluation.

We also selected naturally occurring CREs from the human genome to investigate how well these sequences drive cell type-specific activity compared to our synthetic designs. H3K27ac histone marks and chromatin accessibility as measured by DHS are common proxies for active CREs^6,58^. Thus, for each cell line we identified 4,000 ‘DHS-natural’ sequences with cell type-specific chromatin accessibility and overlapping H3K27ac signals (12,000 total) (**Methods**). We then scanned the entire human genome for 200-mers predicted to be cell type-specific by Malinois and selected 4,000 ‘Malinois-natural’ sequences with the greatest on-target expression and minimal off-target expression in each of the three cell lines (**Methods, Supplementary Figure 8a**). Notably, there was low overlap between elements identified using DHS or Malinois (0.10%-4.1% intersection depending on cell type of interest, **Supplementary Figure 8c**). Although DHS-natural sequences displayed high levels of chromatin accessibility, Malinois-natural and both synthetic groups were predicted to have greater cell type specificity, with non-penalized synthetic sequences surpassing all groups (**Supplementary Figure 9**).

All methods used to generate synthetic CREs resulted in groups of sufficiently diverse sequences. We first quantified single-nucleotide similarity by calculating the average Levenshtein distance of each sequence to its 4 nearest neighbors within the corresponding design group, and repeated this process for human promoters and shuffled sequences from the library as controls (**Supplementary Figure 10a**). DHS-natural, and non-repetitive Malinois-natural sequences were respectively 1.2%, and 11.8% closer to neighbors than shuffled controls. Depending on the generative algorithm, non-penalized synthetic sequences were 0.57%-2.9% closer to neighbors. Interestingly, synthetic-penalized sequences were on average 0.45%-0.89% further away from their 4 nearest neighbors than shuffled controls, with distances increasing during successive penalization rounds (Spearman’s ⍴=0.73 *p*<10^-300^). In contrast, promoters were 8.9% closer to neighbors than shuffled controls, implying that synthetic sequences are substantially more diverse than promoters. As a more stringent assessment of diversity that can capture reuse of individual sequence motifs, we also quantified the average distance of 7-mer content to the 4 nearest neighbors for all oligos. On average, non-repetitive natural sequences selected by DHS and Malinois were 3.0% and 24.4% closer to their nearest neighbors, respectively, than shuffled sequences. Synthetic sequence pairs showed median levels of 7-mer diversity in between groups of natural sequences, being on average 3.6%-7.2% closer to nearest neighbors than shuffled sequences. Motif penalization significantly reduced neighbor closeness from 6.5% to 0.82% relative to shuffled controls (Spearman’s ⍴=0.75, *p*<10^-300^, **Supplementary Figure 10b**). On the other hand, despite the modest reductions compared to shuffle sequences, all groups except Malinois-natural showed less 7-mer similarity than promoters (on average 9.7% closer to nearest neighbors than shuffled sequences), supporting the conclusion the test library provides a diverse collection of CREs for experimental validation.

### CODA successfully generates synthetic CREs with high cell type specificity

We experimentally tested the library of 77,157 natural and synthetic sequences (**Figure 2b**) to determine if machine-guided sequence design could reliably generate biologically functional elements with desired activity. In total, the library included 51,000 synthetic sequences (36,000 standard and 15,000 motif-penalized), 24,000 natural sequences (12,000 DHS-natural and 12,000 Malinois-natural), and 2,157 experimental controls. We quantified activity of an individual CRE as the log2 fold change (log_2_FC) of expression of the reporter gene driven by the CRE compared to a set of negative controls (**Figure 2b,c**). Empirical MPRA measurements of this library and Malinois predictions were well correlated (Pearson’s *r* 0.79-0.91; Spearman’s ⍴ 0.84-0.92; **Supplementary Figure 11**), suggesting Malinois’ predictive accuracy is not limited to natural sequences.

We were able to identify naturally occurring sequences with cell specificity, with Malinois-natural sequences significantly outperforming DHS-natural sequences, suggesting that DHS peaks are a poor predictor of specificity in MPRA. To quantify cell type-specific expression between design groups we used the MinGap score, which is the log_2_FC in the target cell type minus the maximum off-target log_2_FC. Consistent with *a priori* Malinois activity predictions of genomic sequences, DHS-natural sequences in all three cell types performed poorly as cell type-specific CREs compared to natural sequences identified by Malinois (median MinGap difference Malinois-natural vs DHS-natural: K562 2.78, HepG2 1.84, SK-N-SH 0.57; *p*<10^-258^ for all, one-sided Wilcoxon rank-sum test) (**Figure 2d, Supplementary Figures 9 and 12**). These differences in MinGap were primarily driven by weaker on-target activity for DHS-natural sequences compared to Malinois-natural in K562 (median log_2_FC: DHS-natural 2.06, Malinois-natural 4.54) and HepG2 cells (DHS-natural 1.44, Malinois-natural 2.72), while low on-target activity in SK-N-SH in both groups (DHS-natural 0.64, Malinois-natural 0.84) resulted in a lower MinGap difference and reduced SK-N-SH specificity observed in natural sequences in general.

Synthetic sequences from all three algorithms outperformed both groups of natural sequences as cell type-specific CREs in all three cell types. Compared to Malinois-natural, the best performing natural sequence group, synthetics displayed a higher MinGap for all target cell types (median MinGap difference synthetics vs Malinois-natural: K562 1.70, HepG2 0.65, SK-N-SH 2.28; *p*<10^-121^ for all, one-sided Wilcoxon rank-sum test) (**Figure 2d, Supplementary Figure 12**). Performance gains of synthetic sequences were primarily driven by greater repression in off-target cell types (median off-target log_2_FC: synthetic -0.69, Malinois-natural 0.09, DHS-natural 0.41). In addition, synthetic sequences had a higher on-target activity in SK-N-SH (median log_2_FC 3.20) compared to both natural groups, and higher on-target activity for HepG2 and K562 compared to DHS-natural sequences (**Figure 2c**). In summary, synthetic sequences consistently achieved the largest quantitative separation between target and off-target cell types when compared to both classes of naturally derived sequences.

In addition to evaluating specificity using the MinGap, we quantified and visualized specificity utilizing all three cell measurements. We developed a radial coordinate system where the most specific sequences trend outwards along one of the three cell type axes, while sequences with uniform activity across cell types are drawn toward the origin (**Figure 2e, Methods**). The system incorporates both the MinGap and the MaxGap (log_2_FC separation between the target cell type and minimum off-target) scores. We categorize CREs as cell type-specific if two conditions are met: (i) the MaxGap is greater than 1, and (ii) the MinGap:MaxGap ratio is greater than 0.5. These two requirements prioritize sequences with on-target preference while avoiding sequences in which one off-target cell type is closer to the target cell type than the other off-target cell type (**Methods**).

Using our criteria to categorize cell type-specific CREs, we observe that most (94.1%) synthetic sequences designed by CODA successfully drive cell type specificity (**Figure 2e, Supplementary Figure 13**). Depletion of the most optimal motifs did not impact success substantially, with 92.4% of motif-penalized sequences still driving specificity. Comparatively, we observe that Malinois-natural (73.6%) and DHS-natural sequences (40.6%) were less successful (**Figure 2e**). When increasing the stringency of the MaxGap four-fold, synthetic sequences (54.7% specific) further outperformed Malinois-natural (21.5%) and DHS-natural (4.7%) sequences, as well as motif-penalized sequences (30.8%). Overall, synthetic CREs lacking any homology to the human genome (**Methods**) can drive the most consistently robust cell-specific activity in large part through repression of off-target activity, as well as through some increases in on-target activity.

### The TF vocabulary of synthetic sequences drives cell type-specific CRE activity

Having found that synthetic CREs are more cell type-specific than both classes of natural sequences, we sought to link sequence content to the responsible regulatory syntax. Transcription is controlled in part by individual TF binding to sequence motifs as well as interactions between TFs^11^. We identified 82 short (6-15 bp) sequences enriched in our MPRA-tested library, 55 of which can be confidently aligned to a known TF binding motif (**Supplementary Tables 5 and 6**)^66 67^. To interpret the effect of TF binding on sequence function, we predicted single-nucleotide contributions on regulatory activity in each of the three cell types using a robust modified version of Integrated Gradients (**Methods**)^68^.

The regulatory activity contribution scores identify the overall magnitude and direction of the effect of each motif in each of our three cell lines (**Figure 3a**). Of the 82 enriched motifs, 67% had positive predicted contributions to sequence activity while the remaining 33% were repressive. This included well-known activators such as GATA1^69^, a heavily utilized and essential TF expressed in K562, which is correctly predicted by Malinois to drive activity exclusively in K562 (**Figure 3b**). Likewise, HNF1B and HNF4A, master regulators expressed in hepatocyte development^70–73^, are used to drive transcription in HepG2 cells and their contributions are exclusive to HepG2. Motifs displaying negative contributions included the repressors GFI1 in K562^74–76^, and MEIS2 in HepG2 and SK-N-SH^77–79^.

**Figure 3.**
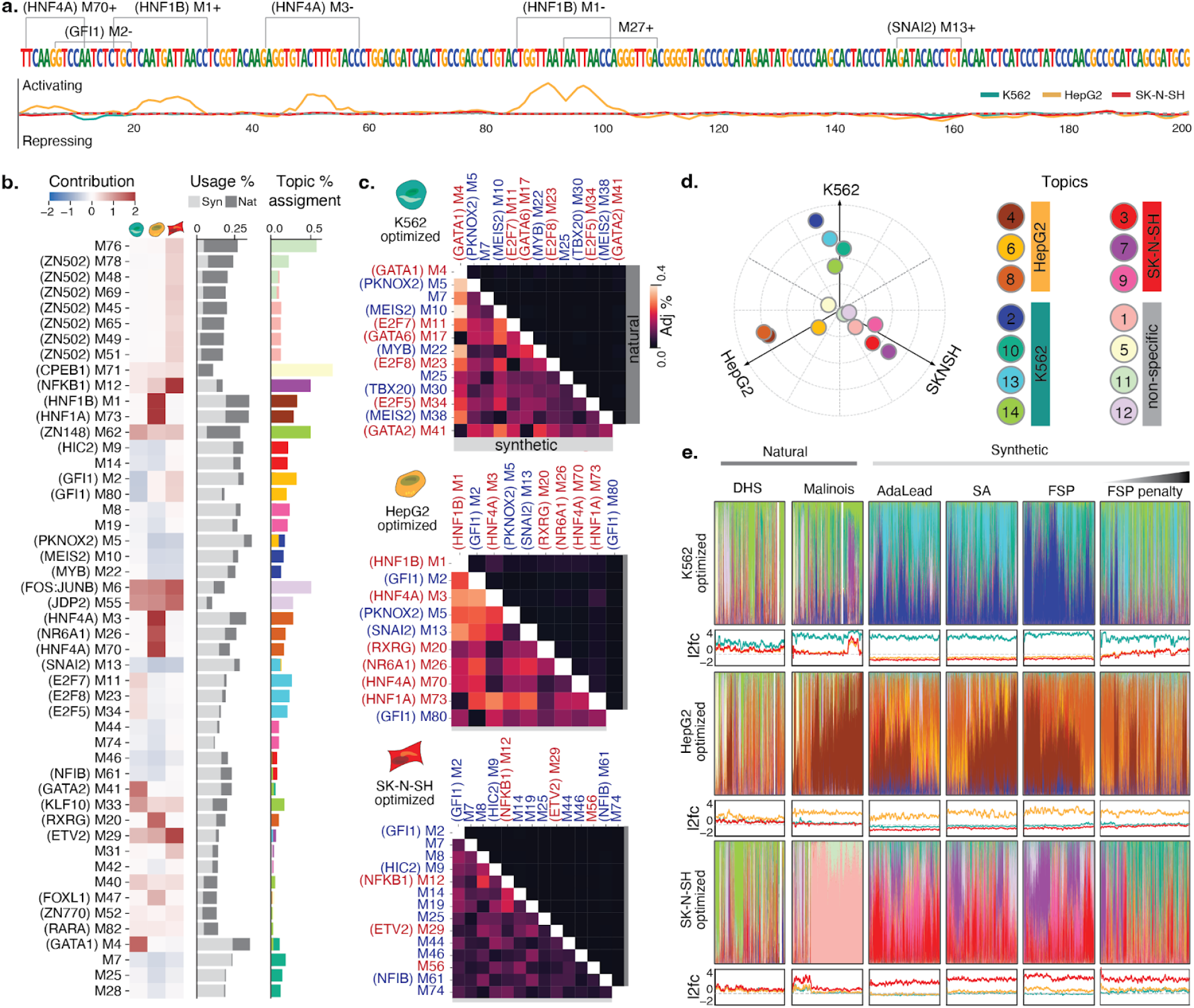
Interpreting CRE syntax in engineered elements. (**a**) Malinois contribution scores enable nucleotide resolution interpretation of sequence activity. Shown is a representative synthetic CRE designed to drive HepG2-specific reporter expression. Enriched motifs, demarcated on the upper sequence track, can be combined with model prediction contribution scores, plotted for each cell type on the lower track (K562: teal, HepG2: yellow, SK-N-SH: red), to interrogate and assign functional subunits. Positive and negative values indicate sequences contribute to transcriptional activation or silencing, respectively, in the corresponding cell type. Motifs are labeled with an “M” followed by their STREME output index. Motifs with a strong known-motif match (Methods) have the name of the match in parenthesis preceding their label. “+” and “-” denote forward and reverse orientations respectively. (**b**) Left heatmap: average contributions of enriched motifs in K562, HepG2, SK-N-SH (left to right columns). Center bar plot: motif enrichment in synthetic (light gray) and natural (dark gray) sequences. The *x*-axis represents the percentage of sequences in each group that contain at least one instance of that motif denoted on the *y*-axis. Right bar plot: motif program association derived from the NMF features matrix. Colors correspond to programs listed in Fig 3e. Only motifs with the top-4 assignments for each topic were included in the figure (see **Supplementary Fig. 14** for full figure). (**c**) Cooccurrences of enriched motifs are more prevalent in synthetic CREs. Adjusted co-occurrence percentage is calculated by multiplying (i) the percentage of sequences in each group containing a pair of motifs and (ii) the similarity divergence of the motifs (1 minus the Pearson correlation coefficient of the motif logos in their optimal alignment) (**Methods**; see **Supplementary Fig. 16** for raw percentages.). Upper and lower triangular percentages correspond to natural and synthetic sequences respectively. Red and blue motif labels denote motifs with mostly positive or negative contribution, respectively. (**d**) Specific functional programs drive cell type-specific transcription. Empirical program function calculated using a weighted average of MPRA log2FC scores based on topic mixture displayed in panel **c**. Ten cell type specificity-driving programs were identified using the same criteria applied to identify cell type-specific sequences (bright colored points; 4 for K562, 3 for HepG2, 3 for SK-N-SH). Four programs are not associated with cell type-specific transcription (pastel points). (**e**) Synthetic and natural sequences show distinct patterns of higher order arrangements of TF binding motifs. Colored bar plots generated from NMF decomposition of synthetic and natural sequences based on enriched motif content reveal the functional programs used in each sequence. For each sequence, programs colored based on the key in **d** and are plotted as a fraction of total program content. Note, in a few cases, sequences were not assigned to any program with any frequency yielding a blank bar. Line plots display MPRA log_2_FC scores for the above sequences in K562 (teal), HepG2 (yellow), and SK-N-SH (red). Sub-panels are organized into rows by expected target cell type and columns by method used to nominate candidate sequences. Sequences in each panel are sorted by hierarchical clustering based on program content.

We examined whether motif use differed between natural and synthetic sequences. All of the 82 enriched motifs occur at least once in both synthetic and natural sequences, suggesting a shared vocabulary between the two classes (**Figure 3b, Supplementary Figure 14 and 15**).

However, the utilization of motifs differed. For example, motifs for transcriptional activators GATA1 in K562 and HNF4A in HepG2 were deployed at higher rates in synthetic sequences (all synthetics: 65.0%, 62.8%, respectively; all naturals: 28.5%, 26.9%, respectively), as well as the repressors MEIS2 in K562 and GFI1 in HepG2 (all synthetics: 58.5%, 51.5%, respectively; all naturals: 5.4%, 5.3%, respectively) (**Supplementary Figure 15**). Overall, fewer motifs were overrepresented (2-fold) in DHS (9/82) and Malinois-natural sequences (18/82) compared to synthetics (38/82).

Notably, we also observed a higher use of particular motif combinations in synthetic sequences that were almost absent in natural sequences. For example, among synthetic sequences, we see higher rates of GATA1/MEIS2 in K562 (46.8%) and HNF4A/GFI1 in HepG2 (31.1%), compared to natural sequences (1.7% and 2.2% each pair respectively) (**Figure 3c, Supplementary Figure 16, Methods**). Across all three cell types, we observed 176 commonly used motif pairs with 100-fold higher utilization in synthetics compared to either DHS or Malinois-natural, and 23 pairs unique to synthetic sequences entirely (**Methods**). Combinations of two distinct activating motifs were observed in most non-penalized synthetic and Malinois-natural sequences (97.1% and 93.6%, respectively), while activating-repressive and repressive-repressive motif pairs were observed at much lower rates in the natural group (activating-repressive: synthetic 98.5%, Malinois-natural 38.3%; repressive-repressive: synthetic 95.5%, Malinois-natural 15.5%), suggesting that natural sequences are less likely to use repressive grammar in constructing cell type-specific CREs.

Further emphasizing the increased use of individual and combinations of motifs in synthetic sequences, we observe that non-penalized synthetic elements showed a greater diversity of unique motifs (types) per sequence (median 2.5-fold vs natural non-repetitive; *p*<10^-300^, one-sided Wilcoxon rank-sum test) as well as a greater number of total motif instances (tokens) (2.3-fold vs natural non-repetitive; *p*<10^-300^, one-sided Wilcoxon rank-sum test) per sequence (**Supplementary Figure 17**). As expected, penalization rounds for synthetic sequences reduce some individual motif instances, reducing both types and tokens (type median 1.5-fold vs natural non-repetitive; token median 1.33-fold vs natural non-repetitive). However, the type:token ratio, a measure of non-redundant motif deployment, is higher in penalized synthetic sequences than in non-penalized ones due to reduced motif redundancy (median type:token 0.78 vs 0.71 respectively; *p*<10^-300^, one-sided Wilcoxon rank-sum test). As these sequences remain highly specific, CODA is able to explore alternative regulatory mechanisms successfully despite increased syntactical constraints posed by penalization.

### Complex semantic architectures are syntactically differentially deployed in natural and synthetic sequences

In addition to single TF-motif usage and pair-wise co-occurrence, cell type specificity is thought to arise through higher-order motif semantics, which can mediate the complex organization of many TFs to impart CRE activity^7,8,11,12^. To aggregate semantically-related enriched motifs into functional programs, we used Non-negative Matrix Factorization (NMF)^80^ to decompose sequences in our library into a mixture of 14 functional programs based on enriched motif content (**Supplementary Figure 18, Methods**). NMF identified 9 programs associated with cell type-specific activity (4 programs in K562, and 3 in each HepG2 and SK-N-SH), with the 4 remaining programs associated with pleiotropic activation and/or repression (**Figure 3d**).

Natural and synthetic sequences deploy semantically distinct programs (**Figure 3e, Supplementary Figure 19**). Notably, average program content in synthetic sequences attributed to cell type-specifying programs was significantly higher (K562 86.7%, HepG2 76.3%, SK-N-SH: 70.5%) than in both DHS-natural sequences (K562 46.5%, HepG2 37.6%, SK-N-SH 18.5%; *p*<10^-300^ for all, two-sided Welch’s *t*-test) and Malinois-natural sequences (K562 60.8%, HepG2 73.4%, SK-N-SH 22.7%; *p*<10^-11^ for all, two-sided Welch’s *t*-test) (**Supplementary Figure 20a**). Despite the increased redundancy of certain motifs (such as GATA1 and the HNFs) in synthetic sequences, these CREs have more program heterogeneity than DHS-genomic CREs for all cell types (*p*<1.20e-7, two-sided Welch’s *t*-test) and Malinois-genomic CREs for HepG2 and SK-N-SH-specific candidates (**Supplementary Figure 20b**; *p*<5.92e-96, two-sided Welch’s *t*-test).

We next observed that distinct semantic combinations of programs deployed by CODA contributed to improved specificity in synthetic CREs. When only considering cell type-specific programs, we found natural sequences primarily rely on activating programs while synthetic sequences additionally utilize programs that generally drive off-target cell type repression (median repressing program content: DHS-natural 0.37%; Malinois-natural 0.34%; synthetic 36.8%) (**Supplementary Figure 20c,d**). A plurality of synthetic sequences (45.3%) are substantially composed of both activating and repressing programs, supported by enhancer and silencer TF motif matches, while relatively few DHS (3.4%) and Malinois (6.4%) natural sequences show this combination (**Methods**; **Supplementary Figure 20e**). These results support our motif-based observations that the improved performance of synthetic sequences is due to a combination of on-target activations and off-target repression.

### Synthetic CREs drive desired tissue-specific activity *in vivo*

To assess specificity of synthetic CREs beyond an episomal reporter context in cell lines, we evaluated selected sequences *in vivo* for their ability to drive cell type-specific expression. Using empirical MPRA results, Malinois contribution scores, *in silico* predictions of tissue-specific epigenetic signals, and element syntax (**Methods, Supplementary Figure 21**), we nominated three liver- and three neuronal-specific CREs for *in vivo* characterization in zebrafish embryos (**Supplementary Figure 22**).

We inserted synthetic sequences upstream of a minimal promoter driving GFP to emulate the vector design utilized by CODA during *in vitro* testing^81^. We injected transposon vectors into embryos and integrated them into the zebrafish genome. To identify the unique expression patterns of each regulatory element, we performed high-resolution, whole-animal imaging at 48 and 96 hours post fertilization for neuronal and liver targets respectively. For sequences designed to drive activity specifically in the liver, 2 of 3 sequences demonstrated strong, consistent expression in developing hepatocytes (**Figure 4a, Supplementary Figures 23 and 24**). Remarkably, we detected minimal off-target expression in non-targeted cell types.

**Figure 4.**
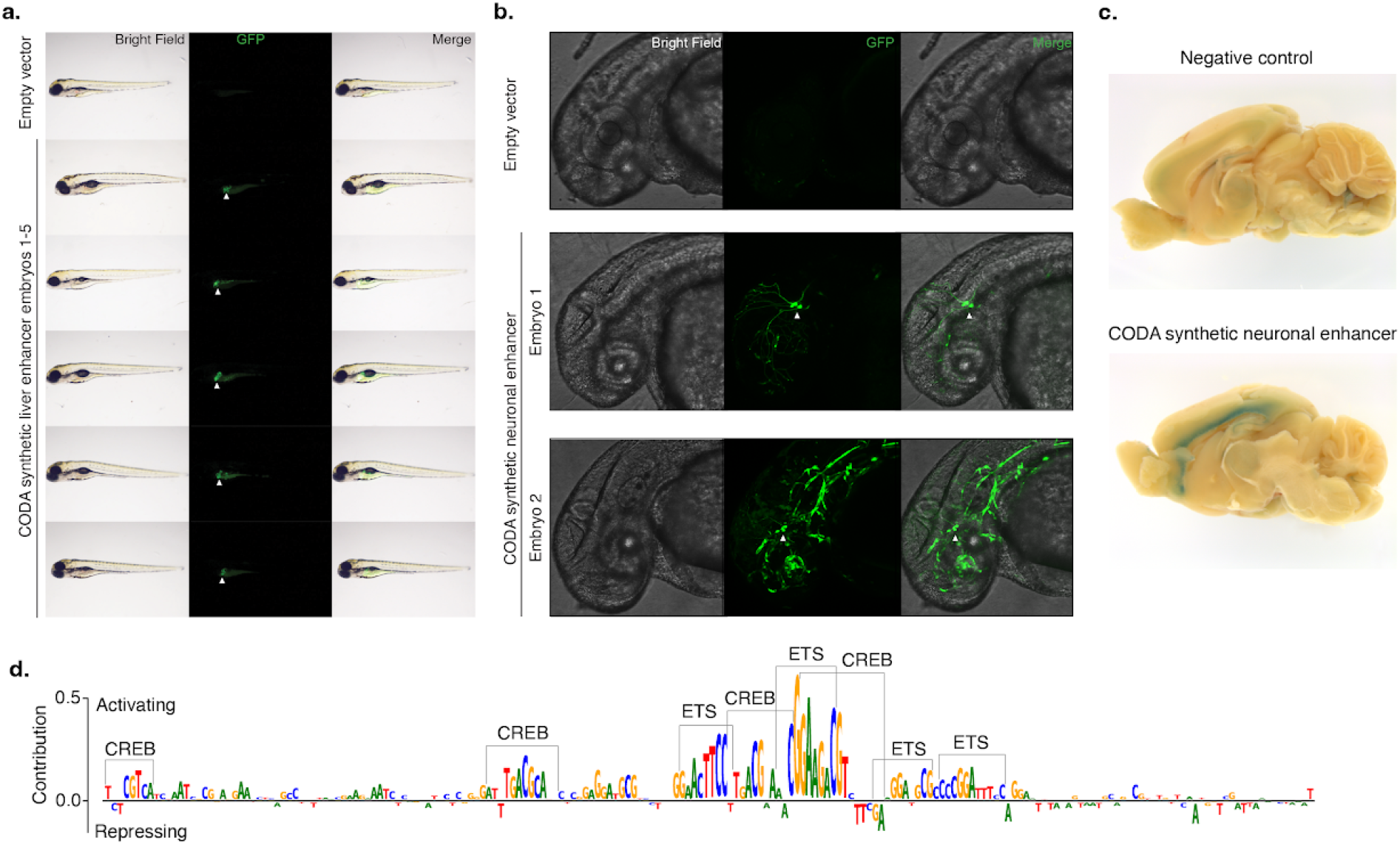
*In vivo* validation of synthetic elements using zebrafish and mouse. (**a**) A synthetic liver-specific CRE drives transgene expression in the larval zebrafish liver. Brightfield, GFP, and merged whole animal imaging 96 hours post-fertilization indicates that the synthetic CRE reproducibly drives transgene expression in zebrafish liver (white arrows). Lateral view, anterior to the left, dorsal up. (**b**) CODA-designed SK-N-SH-specific CRE drives GFP expression in embryonic zebrafish neurons (white arrows). Brightfield, GFP, and merged imaging of the brain and anterior spinal region of animals 48 hours post-fertilization show transgene expression in the developing brain and spinal cord. Embryo 2 shows additional incidental off-target expression in vascular tissue. Lateral view, anterior to the left, dorsal up. (**c**) Synthetic SK-N-SH-specific CRE drives transgene expression in 5-week-old postnatal mice. X-Gal staining for LacZ of the medial section of the brain reveals specific transgene expression at layer 6 of the neocortex. (**d**) Malinois contribution scores reveal the role of ETS and CREB-like binding domains in mediating synthetic CRE activity in neurons. Subsequences of high predicted contribution to SK-N-SH activity overlap with ETS- and CREB-like binding motifs based on visual inspection.

Sequences designed for neuronal specificity showed similar success (2 of 3), driving expression in a subset of neuronal cell types (**Figure 4b, Supplementary Figure 25**). For both successful neuronal-nominated CREs, we observed GFP expression within cell bodies and axonal projections of the developing brain and spinal cord (**Figure 4b, Supplementary Figure 25h**).

We next evaluated if the activity of the two sequences with neuronal specificity in zebrafish extended to a mammalian mouse model system. We placed each synthetic CRE sequence into a targeting vector upstream of a minimal promoter driving lacZ and GFP, and integrated the construct at the H11 safe harbor locus of the mouse through zygote microinjection^82^. We harvested embryos at embryonic day 14.5, a time point roughly equivalent to that used in zebrafish, and used lacZ staining to the transgenic embryos to examine expression patterns of the reporter construct driven by the synthetic CRE. We observed specific expression for neuronal #1 with localized expression in the developing cortex and no additional expression observed elsewhere (**Supplementary Figure 26a**). To localize the expression patterns further within the cortex, we repeated the reporter assay with the neuronal #1 CRE and performed *in situ* staining of the whole brain at 5 weeks postnatal (**Figure 4c, Supplementary Figure 26c-h)**. We confirmed cortex specific expression with focal activity occurring at neocortical layer 6, confirming its neuron-specific activity.

Having designed and validated a novel CRE with strong neuronal specificity, we sought to further elucidate the factors responsible for transcriptional activity in neuronal cells. Using Malinois’ single-nucleotide contributions generated for neuronal sequence #1 in SK-N-SH, we observed two categorically distinct motif classes as contributors to sequence activity: (i) three primary ETS GGA(A/T) binding domains, and (ii) four CREB-like TGACGCA binding domains (**Figure 4d**). ETS factors constitute one of the largest transcription factor families, and its members exhibit highly similar binding motifs. Previous work has reported the potential of ETS factors to form heterodimers with CREB^83^, and our contribution scores provided support for three heterodimer pairings in the sequence. In the off-target cell types, ETS and CREB-like motifs were either reduced or absent, with the presence of two additional negatively contributing motifs, closely matching the repressor GFI1 (**Supplementary Figure 22d**). This suggests that the specificity of neuronal sequence #1 could be partly attributed to the on-target transcriptional activity of cooperative heterodimers and off-target repression by GFI1.

## Discussion

In this study, we developed CODA, an effective strategy to design novel synthetic CREs that can direct cell type-specific gene expression by understanding the complex combinatorial rules of *cis*-regulatory control. CODA builds on previous attempts to design CREs^7,9^, by uniquely combining advances in sequence generation algorithms with the accuracy of Malinois, our CNN model of CRE activity. Synthetic sequences easily outperform natural sequences in driving cell type-specific gene expression, which suggests that novel functions can be programmed into CREs and interpreted by human cells. Using high-throughput characterization methods and *in vivo* reporters, we empirically validated that CODA can design specific CREs with high success rates.

The dearth of natural sequences capable of achieving exquisite cell specificity in our study highlights the difficulty of using human genomic sequences to achieve non-natural objectives for which evolution has not necessarily optimized. Furthermore, DHS elements exhibited both weak on-target activity and poor specificity, possibly a reflection of selective pressure that has shaped DHS elements across mammalian evolution to be optimized for redundancy, versatility, and modular function^84,85^. Without human input, CODA deploys unique combinations of strongly on-target activating and off-target repressing TFs within a short sequence that are not commonly found in the human genome, to yield highly specific synthetic CREs. This suggests that our models have learned a component of the foundational rules governing CREs, and possess the ability to extrapolate this knowledge to unobserved or rarely observed syntax combinations.

Using Malinois, we were able to identify sequences in the genome with moderate proficiency for cell-specific activity, albeit to a lesser degree than synthetics. It was striking that these cell-specific natural sequences represented a broad range of genomic annotations and were less likely to be attributed to known CREs that were found using epigenomic signatures. This highlights the need to carefully consider sequences outside the typically studied candidate CREs when generating libraries with the intent to train high-performance models.

Our high success rate in modeling, generating, and testing sequences *in vitro* prompted us to extend assessment *in vivo*. Despite potential challenges of incomplete conservation of tissue types, heterochrony, and lineage-specific regulatory grammar, our CREs displayed conserved cross-species activity in zebrafish and mice. Our results suggest that CREs designed for tissue-specific targeting can work across species, even in the brain, which has been an ongoing challenge to target with viral-based delivery approaches^41^. An integrated framework leveraging human cell lines in conjunction with whole organism models may thus be a viable approach to rapidly identify CREs to execute novel functions in humans.

We expect that the CODA platform can be extended by integrating additional advancements in deep learning and generative AI, conditioning models on orthogonal data modalities, modeling CRE function in more tissue types, and tasking different biological objectives. While we only tested three cell types here, there is a growing list of clinically actionable tissues that could be benefited, as well as cell types that suffer toxic off-target tropism that could be mitigated by engineered CREs paired with delivery systems. Applying MPRA in additional cell types with greater clinical relevance will enable CODA to better design CREs with specificity tailored for therapeutic applications. While we successfully deployed CODA to maximize cell type specificity, the platform is designed to be flexible to any objective function. We could deploy it to design CREs for drug responsiveness (e.g. glucocorticoids), fine tune expression outputs, or to respond to the complex syntax specific to cancer cells. CODA has improved our ability to write regulatory code tailored to diverse purposes, and could serve as a valuable platform for improving specificity of gene therapies.

## Methods

### Training Malinois, a model of MPRA activity of CREs

To enable systematic evaluation of parameters governing data preprocessing, model architecture, and training we developed tools for limited automatic machine learning in PyTorch (https://github.com/sjgosai/boda2). We implemented support for regression based on DNA sequences using convolutional neural networks. We deployed a containerized application based on this library in conjunction with the Vertex AI platform on Google Cloud to tune all hyperparameters using Bayesian Optimization.

### Data preprocessing

To construct the train/validation/test dataset to train Malinois, we aggregated the log_2_FC output of sequences tested in K562, HepG2, and SK-N-SH from multiple projects. The majority of projects focused on testing the allelic effects of human genetic variation with the remaining projects testing only the reference sequences of the human genome. In total, 776,474 (813,051 before applying filters) unique oligos were aggregated, originating from 10 independent experiments (from three different projects: UKBB, GTEx, BODA). Oligos with a plasmid count less than 20 or no RNA count in any cell type were discarded. The log_2_FC of oligos present in more than one UKBB library was averaged across libraries. If an oligo in UKBB was also found in GTEx or BODA, only the UKBB readout was collected and the others were discarded. If an oligo in GTEx (but not in UKBB) was also found in BODA, only the GTEx readout is collected and the BODA readout was discarded. Non-natural sequences from BODA were discarded.

Also, oligos with a log_2_FC 6 standard deviations below the global mean were discarded (less than 10 oligos). Sequences were padded on both sides with constant sequences from the reporter vector backbone to form 600-bp sequences and converted into one-hot arrays (i.e., A := [1,0,0,0], C := [0,1,0,0], G := [0,0,1,0], T := [0,0,0,1], N := [0,0,0,0]). Oligos from chromosomes 19, 21, and X were held out from the parameter training loop as a validation set guide hyperparameter tuning. Oligos from chromosomes 7, 13 were held out from both parameter training and hyperparameter tuning loops as a test set for reporting performance. Data augmentation was performed by including into the training set the reverse complement of the (600-bp) sequences, and duplicating oligos that had a log_2_FC greater than 0.5 in any cell type.

### Model architecture

The final Malinois model is composed of three functional segments: (1) three convolutional layers with batch normalization and maximum value pooling, (2) a linear layer to integrate positional and feature information from the previous layers, and (3) a stack of branched linear layers such that each output feature is a function of 4 independent linear transformations. As the first two segments are based on the Bassett architecture^46^, Malinois accepts batches of 4 x 600 arrays corresponding to one-hot encoded DNA sequences, so predictions are made by padding inputs on both sides with constant sequences from the reporter vector backbone.

### Model fitting

We trained Malinois using the Vertex AI API on the Google Cloud Platform (GCP). This enabled optimization of all tunable parameters controlling data preprocessing, model architecture, and model training. To do this, first we generated a docker container (gcr.io/sabeti-encode/boda/production:0.0.11) with an installation of CODA using a GCP VM with the following specifications: Debian based Deep Learning VM for Pytorch CPU/GPU operating system, a2-highgpu-1g machine type, and 1 NVIDIA Tesla A100 40G GPU. The container entrypoint was set to a python script for model training (boda2/src/main.py). Using this container we deployed Hyperparameter Tuning Jobs using the default algorithm to optimize the indicated hyperparameters (**Supplementary Table 7**).

### Correlation of Empirical and Predicted MPRA Activity

When comparing Malinois’ predictions to empirical MPRA, we discard any oligo with a replicate log_2_FC standard error greater than 1 in any cell type. Malinois’ predictions for the (padded) forward and reverse complement sequences are averaged into a single prediction.

### Optimization of Cell Specificity

The objective function to guide the sequence design with Simulated Annealing (minimize energy) was the MinGap (Malinois log_2_FC prediction in the target cell type minus the maximum off-target cell type log_2_FC prediction). The objective function used with the algorithms Fast SeqProp and AdaLead (minimize loss or maximize fitness respectively) was the bent-MinGap, which is defined as follows. Let *y*_+_ be the Malinois log_2_FC prediction on the target cell type, and *y*_-_ the maximum of the log_2_FC predictions on the off-target cell types of a given sequence (so MinGap = *y*_+_ - *y*_-_). We constructed a bending function *g*(x) = *x* - *e^-x^* + 1 to preprocess predictions such that the objective function becomes bent-MinGap = *g*(*y*_+_) - *g*(*y*_-_). We applied *g*(*x*) to the predictions to incentivize greater MinGaps with low expression in the off-target cell types. For three generative algorithms, Malinois predictions were clippled to an interval (default: [-2, 6]) to avoid prioritizing sequences with pathologically unrealistic log2FC activity predictions.

### Iterative Maximization of Sequence Fitness Using Iterative, Generative, and Evolutionary Sequence Generation Algorithms

#### Fast SeqProp^36^

We implemented this algorithm as described in previous work but we removed the learnable affine transformation in the instance normalization layer and drew many one-hot encoded samples from the categorical nucleotide probability distribution in each optimization step to more confidently estimate the gradients of the learnable re-parameterized input sequence. The input parameters were randomly initialized (drawn from a normal distribution) and optimized using the Pytorch implementation of the Adam optimization algorithm with a learning rate of 0.5, along with a Cosine Annealing scheduler with a minimum learning rate of 10^-6^ over 300 training steps. In each training step, the loss function value was the negative average bent-MinGap of 20 sequence samples drawn from the categorical nucleotide probability distribution at that step. Once optimization is finalized, instance normalization is applied to the learned input and 20 sequences were sampled from the obtained distribution, and the sequence with the highest predicted bent-MinGap was collected unless the value was less than 3.6.

#### AdaLead^37^

We implemented this algorithm as written in the GitHub repository associated with the original paper. In each run, 20 randomly initialized sequences are optimized over 30 generations with mu=1, recomb_rate=0.1, threshold=0.25, rho=2, using bent-MinGap as the fitness function. Once optimization is finalized, only the sequence with the highest predicted bent-MinGap is collected unless the MinGap was less than 2. We chose to collect only one sequence per run to maximize diversity in the global batch collected from all runs.

#### Simulated Annealing^65^

We implemented Simulated Annealing based on the Metropolis-Hastings algorithm for Markov Chain Monte Carlo simulations. Proposals were generated symmetrically at each step by mutating 3 random bases. We used negative MinGap (without bending) to simulate the energy landscape of the theoretical system. During optimization the temperature term was reduced using a monotonically decreasing function with a diverging infinite sum. (INSERT EQN temp = 1 / ((1+step)^0.501)). To produce sequences with high target-specific activity we used negative MinGap (without bending) to simulate energy of the system.

### Motif Penalization

For each target cell type, the iterative sequence generation penalizing motifs from previous rounds was done in 4 “tracks” (a total of 12 penalization tracks). Each penalization track generates a total of 1750 sequences as follows. First, a batch of 500 sequences (round 0) is generated free of any motif penalty in the objective function besides target cell specificity. Then, round-0 sequences are analyzed for motif enrichment (10 motifs of length 8 to 15) using PyMemeSuite, and the top motif from the enrichment output list is added to a pool of unwelcome motifs in forward and reverse complement orientations. The next batch of 250 sequences to be generated (round 1) has an additional term in its objective function that penalizes the presence of motifs from the pool of unwelcome motifs. This process iterates to complete 5 penalization rounds, 6 rounds in total (500 unpenalized sequences and 5*250 penalized sequences). The motif penalty is calculated using the PWMs (log probabilities) of the motifs as filters of a convolutional layer that scans and scores a batch of sequences. The sums of motif scores above a certain motif threshold in each sequence are averaged and divided by the batch size and the number of filters in the convolutional layer. Each motif threshold is calculated as score_pct * max_score, where max_score is the PWM score of the motif consensus sequence and score_pct is a scaling parameter (0 for K562, 0.25 for HepG2 and SK-N-SH). We also included a penalty weight to each motif in the pool to slightly emphasize the penalty of motifs from earlier rounds as the pool increases. The penalty weight is defined as (current_round_index - motif_round_index)^(1/3) where motif_round_index is the index of the round from which a motif was extracted and added to the pool. Each penalty weight scales the PWM of its respective motif.

In **Supplementary Figure 7b**, the motif-presence score (*y*-axis) of a motif in each sequence was calculated by summing all the motif-match scores that pass the patser score threshold as defined in Biopython86, and then dividing by the maximum possible motif score (the match score of the motif consensus sequence).

### Homology search using Nucleotide Blast

We conducted homology search using NCBI ElasticBLAST to determine if synthetic sequences had measurable homology the any sequences in Nucleotide Collection. We used the blastn algorithm, the dc-megablast task, and a word size of 11 and maintained the defaults for all other settings.

### Selection of Naturally Occurring Cell-Specific Sequences by DNase and Malinois Driven GenomeScan

#### DHS-natural

To identify CREs broadly replicating across experimental approaches, we first took DNAse peaks from each of the three cell lines (K562, HepG2, and SK-N-SH), and subsetted peaks that intersected with H3K27ac peaks from the same cell type. For the DHS-H3K27ac peaks, in each cell type, we scored the average K562, HepG2, and SK-N-SH DHS signal in the peak. We then calculated the MinGap score for each target cell type using the DHS signal, and selected the 4000 peaks with the largest MinGap score in each cell type.

#### Malinois-natural

To nominate cell-specific natural sequences with Malinois, we tiled the whole human genome into 200-bp windows using a 50-bp stride and generated predictions for each window sequence. The cell specificity fitness of each sequence was obtained by evaluating the fitness function mentioned above (bent-MinGap), and the top 4000 best performing sequences were selected for each cell type.

### Genome Annotation of Natural Sequences

Malinois-natural sequences capture a unique component of the genome compared to DHS-natural, with 2.7% of Malinois-natural sequences overlapping sequences in our DHS-natural set, and 65.8% residing outside any previously annotated CREs. cCRE BED files for promoter-like sequences, proximal enhancer-like sequences, distal enhancer-like sequences, and CTCF-only were downloaded from the ENCODE SCREEN Portal^5^ and concatenated into a single BED file for intersection with DHS-natural and Malinois-natural BED files using a custom script. Intersections were done with bedtools 2.30.0^87^ and pybedtools 0.9.0^88^ with the following command ‘Malinois/DHS-natural_BED.intersect(ENCODE_cCRE_BED, wa=True, u=True) and the number of intersections were reported. To determine the genomic features overlapping DHS-natural and Malinois-natural sequences, the same BED files were used as input for ‘annotatePeaks.pl from the homer suite v4.11^89^ with the following command ‘annotatePeaks.pl inputBED hg38 -annStats annStats.txt > annotatePeaksOut.txt’. Annotations for the whole genome (hg38) were generated by dividing the genome into 200-bp intervals using the bedtools makewindows command ‘bedtools makewindows -g hg38.txt -w 200 > hg38_200bp.bed’. Annotations were generated for each cell type (K562, HepG2, SK-N-SH) and sequence selection method (DHS-natural, Malinois-natural.)

### Sampled Integrated Gradients to compute contribution scores of Malinois predictions

We calculated nucleotide contribution scores for each sequence in the proposed library using an adaptation of the input attribution method Integrated Gradients^68^. Sampled Integrated Gradients considers the expected gradients along the linear path in log-probability space from the background distribution to the distribution that samples the input sequence almost surely. In each point of the linear path, a sequence probability distribution (a.k.a. Position Probability Matrix) is obtained from the log-probability space parameters by applying the Softmax function along the nucleotide axis, and a batch of sequences is sampled from such a distribution to be fed into the model. We then calculate the gradients of the batch model predictions with respect to the parameters in the log-probability space, using the straight-through estimator to backpropagate through the sampling operation. The batch gradients are averaged for each point in the path and approximate the gradient integral as in the original formulation of the method. In our case, the subtraction of the baseline input from the input of interest involves the parameters in log-probability space.

### Propeller plots

A propeller dot plot (top row of **Figure 2e**) is a 2-dimensional plot scheme of our own device which seeks to elucidate the cross-dimensional non-uniformity of 3-dimensional points. In this coordinate system, a point’s radial distance from the origin corresponds to the difference between the maximum and minimum values. Its deviant angle from the axis corresponding to the maximum value quantifies the position of the median value within the range of the minimum and maximum values. Namely, the angle is proportional to the ratio between two differences: (i) the difference of the median and minimum values, and (ii) the difference of the maximum and minimum values. This ratio represents the 60-degree-angle fraction deviating from the axis corresponding to the maximum value towards the axis corresponding to the median value. A higher angle of deviation (maximum of 60 degrees) indicates that the median value is closer to the maximum value, while a lower angle (minimum of 0 degrees) of deviation indicates that the median value is closer to the minimum value.

This can also be formulated in terms of the MinGap (maximum - median) and MaxGap (maximum - minimum). In our coordinate system, the MaxGap corresponds to the radial distance. The difference (1 - MinGap/MaxGap) corresponds to the 60-degree-angle fraction deviating from the axis corresponding to the maximum value towards the axis corresponding to the median value. The MinGap:MaxGap ratio controls how much a point gravitates toward a main axis and away from the in-between-axis areas. A ratio of 0 means that the MinGap is zero and therefore the median value is equal to the maximum, so the point will be exactly between two axes. If the ratio is 1, it means that the median and the minimum values are equal, therefore the point will fall exactly in the axis corresponding to the maximum value. Note that, in order for this point of view to work with target and off-target cell type activities, we assume that the maximum cell type activity is the intended target cell type. This implies that, when counting sequences that pass specificity thresholds in Figure 2e, some sequences get their target cell type reassigned to the cell type with the maximum activity, with DHS-natural sequences being the group that most benefits from the reassignment. A total of 652 sequences pass the lenient specificity threshold of MaxGap > 1 and MinGap/MaxGap > 0.5 by getting their target cell type reassigned (DHS-natural: 565, Malinois-natural: 39, AdaLead: 12, Simulated Annealing: 5, Fast SeqProp: 0, Fast SeqProp penalized: 4). However, only 16 sequences pass the stringent specificity threshold of MaxGap > 4 and MinGap/MaxGap > 0.5 by getting their target cell type reassigned (DHS-natural: 15, Malinois-natural: 0, AdaLead: 1, Simulated Annealing: 0, Fast SeqProp: 0, Fast SeqProp penalized: 0).

As an example of coordinate calculation, take the point (5, 3, 1). This point would have a radial distance of 5-1=4 and an angle of deviation from the axis of the first dimension of (3-1)/(5-1)*(60 deg) = 30 deg (in the direction of the axis of the second dimension). In terms of the MinGap:MaxGap ratio, the angle of deviation from the axis of the first dimension (the dimension of the maximum value) towards the axis of the second dimension would be (1 - (5-3)/(5-1))(60 deg) = 30 deg. Observe that all the points of the form (x+4, x+2, x), for any real value of x, will have the same coordinates as the point (5, 3, 1).

A propeller count plot (bottom row of **Figure 2e**) shows the percentage of points that fall in each given area of a propeller dot plot. The teal, yellow, and red regions capture sequences in which the median value is closer to the minimum value than to the maximum value.

The two synthetic groups in **Figure 2e** were randomly subsampled to have exactly 12,000 sequences each and avoid over-plotting compared to the plots of the two natural groups. **Supplementary Figure 13** shows the complete propeller plots broken down by design method.

Oligos with a replicate log_2_FC standard error greater than 1 in any cell type were omitted from the plots.

### Motif enrichment analysis

We submitted our entire library (including controls) to STREME^90,91^ for motif enrichment analysis with the default settings: *minimum width* = 8, *maximum width* = 15, *p-value threshold* = 0.05, and shuffled input sequences as control sequences. Then, the 82 enriched motifs (**Supplementary Table 8**) were submitted to TOMTOM^90,92^ to find matches to known motifs in JASPAR CORE (2022)^66^ and HOCOMOCO Human (v11 FULL)^67^. The match with lowest E-value was collected as the top match. Matches with an *E-value* ≥ 0.1 were discarded. We used RSAT tools^93^ to convert the MEME output file into JASPAR format for processing and parsing with Biopython^86^. We observed that the enrichment of these 82 motifs in a given individual design group yielded similar local results to the enrichment of motifs obtained from such a group submitted alone (each group has at least 12,000 sequences), suggesting that there was minimal or no loss of information in the global STREME analysis from the point of view of its algorithm. We submitted the list of enriched motifs and our sequence library to FIMO^94^ to find all the significant motif instances.

### Motif co-occurrence

We say a pair of motifs co-occur whenever a sequence has at least one significant instance (obtained through FIMO) of each motif. By co-occurrence percentage of a motif pair we mean the percentage of sequences in a given group in which the motif pair co-occurs. Adjusted co-occurrence percentage is defined as the co-occurrence percentage of a motif pair times a motif-pair divergence coefficient (0 if the motifs are identical, 1 if the motifs are as un-correlated as possible), where the motif-pair divergence coefficient is defined 1 minus the Pearson correlation coefficient between the two motif logos (Information Content Matrices) in their optimal alignment. Selection of top co-occurrences to display in figures was based on capturing the top 25 adjusted co-occurrence percentages for each cell type. When finding motif pairs with 100-fold higher utilization in synthetics compared to either DHS or Malinois-natural, and 23 pairs unique to synthetic sequences entirely, we require that the motif pairs co-occur in a minimum of 200 sequences.

### Non-negative Matrix Factorization

We used non-negative matrix factorization (NMF) to model semantic relationships between motifs in our sequence library (scikit-learn version 1.2.2, initialized with NNDSVD, Frobenius loss). First we counted enriched motif matches in each sequence with FIMO^94^ to generate X \in N_0^{n x f} where rows represent sequences in the library. The sample matrix X can then be decomposed into the coefficients and features matrices W \in R^{n x k} and H^{k x f), respectively. We tested k \in [8,28] using bi-cross-validation^95^ and identified an “elbow” in the reconstruction error at k=14. For comparative analysis, we normalize the coefficient matrix to sum to 1.

To improve interpretability of the topic modeling, we generated an additional 4000 sequences for each cell type which prioritized off-target expression. To produce these sequences, we clipped predictions outside the [-2,3] range and applied an alternate fitness function. We optimized sequences to minimize the squared distance from a predicted activity of 3 in the on-target cell and an activity of -2 in each off-target cell. These 12000 additional sequences were included in the final NMF decomposition.

### CODA MPRA

#### MPRA library construction

CODA MPRA library was constructed following protocols previously described in Tewhey et al. 2016^14^. In brief, oligos were synthesized (Twist Bioscience) as 230 bp sequences containing 200 bp of genomic sequences and 15 bp of adaptor sequence on either end. The oligo library was PCR amplified with primers MPRA_v3_F and MPRA_v3_20I_R to add unique 20 bp barcodes along with arms for Gibson assembly into a backbone vector. The oligonucleotide library was assembled into pMPRAv3:Δluc:ΔxbaI (Addgene plasmid #109035) and expanded by electroporation into *E.coli*. Seven of the ten expanded cultures were purified using Qiagen Plasmid Plus Midi Kit to reach 200-300 colony-forming units (barcodes) per oligonucleotide. The expanded plasmid library was sequenced on an Illumina NovaSeq using 2×150 bp chemistry to acquire oligo-barcode pairings. The library underwent AsiSI restriction digestion, and GFP with a minimal promoter amplified from pMPRAv3:minP-GFP (Addgene plasmid #109036) using primers MPRA_v3_GFP_Fusion_F and MPRA_v3_GFP_Fusion_R was inserted by Gibson assembly resulting in the 200 bp oligo sequence positioned directly upstream of the promoter and the 20 bp barcode falling in the 3’ UTR of GFP. Finally, the library was expanded within *E.coli* and purified using the Qiagen Plasmid Plus Giga Kit.

#### MPRA library transfection into cells

Two hundred million cells were transfected using the Neon Transfection System 100ul Kit with 5ug or 10ug of the MPRA library per ten million cells. Cells were harvested 24 hours post transfection, rinsed with PBS and collected by centrifugation. After adding RLT buffer (Rneasy Maxi kit), dithiothreitol and homogenization, cell pellets were frozen at -80°C until further processing. For each cell type, 3 biological replicates performed on different days.

#### RNA isolation and MPRA RNA library generation

RNA was extracted from frozen cell homogenates using the Qiagen RNeasy Maxi kit. Following DNase treatment, a mixture of 3 GFP-specific biotinylated primers were used to capture GFP transcripts using Sera Mag Beads (Fisher Scientific). After a second round of DNase treatment, cDNA was synthesized using SuperScript III (Life Technologies) and GFP mRNA abundance was quantified by qPCR to determine the cycle at which linear amplification begins for each replicate. Replicates were diluted to approximately the same concentration based on the qPCR results, and first round PCR (8 or 9 cycles) with primers MPRA_Illumina_GFP_F_v2 and Ilmn_P5_1stPCR_v2 were used to amplify barcodes associated with GFP mRNA sequences for each replicate. A second round of PCR (6 cycles) was used to add Illumina sequencing adaptors to the replicates. The resulting Illumina indexed MPRA barcode libraries were sequenced on an Illumina NovaSeq using 1×20bp chemistry.

### CRE prioritization for *In vivo* validation

#### Enformer analysis of epigenetic signatures

To simulate epigenetic signatures *in silico* we collected the nucleotide sequence from chr11:3,101,137-3,493,091 of the mouse reference genome (mm10). The expected insertion sequence using an H11 targeting vector with a lacZ:P2A:GFP open reading frame was added. As a control, the expected CRE insertion site was simulated as a 200 nucleotide sequence of N. We simulated all possible CRE insertions corresponding to our cell type-specific MPRA by replacing the oligo-N sequence with 200-mers from our library. We inferred epigenetic signatures for all of these sequences using Enformer by modifying the notebook provided by this link (https://colab.research.google.com/github/deepmind/deepmind_research/blob/master/enformer/enformer-usage.ipynb). To estimate CRE induced transcriptional activation in various tissues we collected 128 nucleotide resolution epigenetic signatures overlapping the expected insertion (35 bins). To calculate an aggregate effect for each tissue, we calculated the max signal for each feature over the insertion, followed by a feature-specific Yeo-Johnson power transformation.

Normalized features were then selected based on tissue correspondence (**Supplementary table 8**) and averaged to estimate CRE activity in 10 different tissues.

#### Manual sequence prioritization

Sequences were prioritized based on review of empirical MPRA measurements, contribution scores, motif matches, sequence content, and predicted epigenetic signatures. We looked for sequences that displayed a high separation between the MPRA measures of the target and the off-target cell types. We also looked to capture variations of combinations of motif matches, and we used the contribution scores to visually examine the motif matches and other potentially important sequence content. Finally, we selected sequences with at least moderate tissue specificity in predicted epigenetic signatures.

### Transgenics

#### Transient zebrafish synthetic enhancer assay

To build the synthetic enhancer eGFP reporter, double-stranded oligonucleotides corresponding to synthetic enhancers (200 bp) were synthesized by IDT (GeneBlock). Synthetic enhancers were amplified by PCR with primers that included homology to the plasmid vector E1b-GFP-Tol2 (Addgene plasmid #37845)^81^ and were cloned upstream of the minimal promoter (E1b) to generate the synthetic enhancer eGFP plasmid reporter (pTol2-synthetic enhancer-E1b-eGFP-Tol2) using HiFi DNA Assembly following manufacturer’s instructions (New England Biolabs). Reporter plasmid sequences were verified by Sanger sequencing. To transiently express the synthetic enhancer reporter in zebrafish, plasmids were co-injected with tol2 transposase mRNA into 1-cell stage zebrafish embryos following established methods^96^. Injected embryos were imaged at the indicated days (2 or 4 days-post-fertilization) either by dissecting (Olympus) or confocal fluorescence (Leica SP6) microscope. All zebrafish procedures were approved by the Yale University Institutional Animal Care and Use Committee (IACUC) (Protocol Number 2022-20274).

#### Mouse transgenic reporter assay

An H11 targeting vector with an lacZ:P2A:GFP open reading frame was linearized using PCR containing 2 ng of template, 1 μl of KOD Xtreme Hot Start DNA Polymerase (Sigma 71975), 25 μl of Xtreme buffer, and 0.5 μM forward and reverse primers (H11_bxb_lacZ:GFP_lin_F, pGL_minP_GFP_R; **Supplementary Table 9**) cycled with the following conditions: 94°C for 2 min, 20 cycles of 98°C for 10 s, 56°C for 30 s, and 68°C for 13 min, and then 68°C for 5 min. Amplified fragments were treated with 0.5 uL of DpnI (NEB, R0176S) for 30 min at 37°C, purified using 1× volume of AMPure XP (Beckman Coulter, A63881) and eluted with water. Double-stranded oligonucleotides corresponding to synthetic enhancers with gibson arms were synthesized by IDT (GeneBlock) and assembled into targeting vector using 5 μl of NEBuilder HiFi DNA Assembly Master Mix (NEB, E2621S), 36 ng of linearized vector, and 10 ng of the synthesized fragment in 20 μl total volume for 45 min at 50°C. Transgenic mice were created following the enSERT protocol^82^. A mixture of 20 ng/μl Cas9 protein (IDT 1074181), 50 ng/μl single guide RNA (sgRNA_H11lacZ; **Supplementary Table 9**), 25 ng/μl donor plasmid, 10 mM Tris, pH 7.5, and 0.1 mM EDTA was injected into pronuclear of FBV zygotes. The whole embryo at E14.5 or isolated brain at 5 weeks postnatal were fixed at 4°C for 1 hour in PBS supplemented with 2% paraformaldehyde, 0.2% glutaraldehyde, and 0.2% IGEPAL CA-630. After washing with PBS, the embryos were stained at 37°C overnight in a solution in PBS supplemented with 0.5 mg/ml X-gal (Sigma, B4252), 5 mM potassium hexacyanoferrate(II) trihydrate, 5 mM potassium hexacyanoferrate(III), 2 mM MgCl2, and 0.2% IGEPAL CA-630. The images were taken using Leica M165 for embryos or Leica M125 for brains. All mouse procedures were performed in accordance with the National Institutes of Health Guide for the Care and Use of Laboratory Animals, and were approved by the Institutional Animal Care and Use Committees of The Jackson Laboratory (protocol number 18038).

## Data availability

Reference data sets used in this study are linked and annotated in **Supplementary Table 1**. Processed MPRA data used to train Malinois is available in **Supplementary Table 2**. Processed MPRA data and Malinois predictions for the cell type-specific CRE library designed for this study are available in **Supplementary Table 10**.

## Code availability

CODA is available at https://github.com/sjgosai/boda2.

## Acknowledgements

We thank The Jackson Laboratory Genome Technologies, Genetic Engineering Technologies and Microscopy Core for experimental support; Kevin Peterson for reagents and technical advice; and Taneli Helenius, James Xue, Michael Stitzel, Stephen Rong, Dylan Kotliar, Trevor Sorrells, Thanh Thanh Nguyen, Hannah Dewey, Niketa Nerurkar, Mary Teena Joy, Mackenzie Noon, and Arya Rao for suggestions and conversations about the manuscript. This work was supported by Howard Hughes Medical Institute and by US National Institutes of Health grants UM1HG009435, R00HG010669, and R35HG011329.

## Author Contributions

SJG initiated the model-guided sequence design framework. SJG, RIC, SKR, and RT developed the full study and designed experiments. SJG and RIC developed the CODA software library, Malinois model, and produced the synthetic sequences. NF, SK, and SKR conducted in vitro experiments. RRN and KM conducted in vivo experiments. SJG, RIC, NF, JCB, SKR, and RT performed data analysis. SJG, RIC, JCB, PCS, SKR, and RT interpreted results and drafted the manuscript. PCS, SKR and RT secured funding and supervised the study. All of the authors revised the manuscript and accepted its final version.

## References

1. Wittkopp, P. J. & Kalay, G. Cis-regulatory elements: molecular mechanisms and evolutionary processes underlying divergence. Nat. Rev. Genet. 13, 59–69 (2011).

2. Gasperini, M., Tome, J. M. & Shendure, J. Towards a comprehensive catalogue of validated and target-linked human enhancers. Nat. Rev. Genet. 21, 292–310 (2020).

3. de Boer, C. G. & Taipale, J. Hold out the genome: A roadmap to solving the cis-regulatory code. bioRxiv 2023.04.20.537701 (2023) doi:10.1101/2023.04.20.537701.

4. Heinz, S., Romanoski, C. E., Benner, C. & Glass, C. K. The selection and function of cell type-specific enhancers. Nat. Rev. Mol. Cell Biol. 16, 144–154 (2015).

5. ENCODE Project Consortium et al. Expanded encyclopaedias of DNA elements in the human and mouse genomes. Nature 583, 699–710 (2020).

6. Meuleman, W. et al. Index and biological spectrum of human DNase I hypersensitive sites. Nature 584, 244–251 (2020).

7. Donohue, L. K. H. et al. A cis-regulatory lexicon of DNA motif combinations mediating cell-type-specific gene regulation. Cell Genom 2, (2022).

8. Levo, M. & Segal, E. In pursuit of design principles of regulatory sequences. Nat. Rev. Genet. 15, 453–468 (2014).

9. Taskiran, I. I., Spanier, K. I., Christiaens, V., Mauduit, D. & Aerts, S. Cell type directed design of synthetic enhancers. bioRxiv 2022.07.26.501466 (2022) doi:10.1101/2022.07.26.501466.

10. Avsec, Ž., et al. Base-resolution models of transcription-factor binding reveal soft motif syntax. Nat. Genet. 53, 354–366 (2021).

11. Lambert, S. A. et al. The Human Transcription Factors. Cell 172, 650–665 (2018).

12. Kim, D. S. et al. The dynamic, combinatorial cis-regulatory lexicon of epidermal differentiation. Nat. Genet. 53, 1564–1576 (2021).

13. Shrikumar, A., Greenside, P. & Kundaje, A. Learning Important Features Through Propagating Activation Differences. in Proceedings of the 34th International Conference on Machine Learning (eds. Precup, D. & Teh, Y. W.) vol. 70 3145–3153 (PMLR, 06--11 Aug 2017).

14. Tewhey, R. et al. Direct Identification of Hundreds of Expression-Modulating Variants using a Multiplexed Reporter Assay. Cell 165, 1519–1529 (2016).

15. Ulirsch, J. C. et al. Systematic Functional Dissection of Common Genetic Variation Affecting Red Blood Cell Traits. Cell 165, 1530–1545 (2016).

16. Ernst, J. et al. Genome-scale high-resolution mapping of activating and repressive nucleotides in regulatory regions. Nat. Biotechnol. 34, 1180–1190 (2016).

17. Melnikov, A. et al. Systematic dissection and optimization of inducible enhancers in human cells using a massively parallel reporter assay. Nat. Biotechnol. 30, 271–277 (2012).

18. Klein, J. C. et al. A systematic evaluation of the design and context dependencies of massively parallel reporter assays. Nat. Methods 17, 1083–1091 (2020).

19. Lawler, A. J. et al. Machine learning sequence prioritization for cell type-specific enhancer design. Elife 11, (2022).

20. Movva, R. et al. Deciphering regulatory DNA sequences and noncoding genetic variants using neural network models of massively parallel reporter assays. PLoS One 14, e0218073 (2019).

21. Vaishnav, E. D. et al. The evolution, evolvability and engineering of gene regulatory DNA. Nature 603, 455–463 (2022).

22. Agarwal, V., et al. Massively parallel characterization of transcriptional regulatory elements in three diverse human cell types. bioRxiv (2023) doi:10.1101/2023.03.05.531189.

23. Xue, J. R. et al. The functional and evolutionary impacts of human-specific deletions in conserved elements. Science 380, eabn2253 (2023).

24. Siraj, L. & Ulirsch, J. Functional dissection of complex and molecular trait variants at single nucleotide resolution. In Preparation (2023).

25. Rosenberg, A. B., Patwardhan, R. P., Shendure, J. & Seelig, G. Learning the sequence determinants of alternative splicing from millions of random sequences. Cell 163, 698–711 (2015).

26. Bogard, N., Linder, J., Rosenberg, A. B. & Seelig, G. A Deep Neural Network for Predicting and Engineering Alternative Polyadenylation. Cell 178, 91–106.e23 (2019).

27. Sample, P. J. et al. Human 5′ UTR design and variant effect prediction from a massively parallel translation assay. Nat. Biotechnol. 37, 803–809 (2019).

28. Kelley, D. R. et al. Sequential regulatory activity prediction across chromosomes with convolutional neural networks. Genome Res. 28, 739–750 (2018).

29. Zhou, J. & Troyanskaya, O. G. Predicting effects of noncoding variants with deep learning-based sequence model. Nat. Methods 12, 931–934 (2015).

30. Quang, D. & Xie, X. DanQ: a hybrid convolutional and recurrent deep neural network for quantifying the function of DNA sequences. Nucleic Acids Res. 44, e107 (2016).

31. Jaganathan, K. et al. Predicting Splicing from Primary Sequence with Deep Learning. Cell 176, 535–548.e24 (2019).

32. de Almeida, B. P., Reiter, F., Pagani, M. & Stark, A. DeepSTARR predicts enhancer activity from DNA sequence and enables the de novo design of synthetic enhancers. Nat. Genet. 54, 613–624 (2022).

33. Penzar, D. et al. LegNet: a best-in-class deep learning model for short DNA regulatory regions. Bioinformatics 39, (2023).

34. Avsec, Ž., et al. Effective gene expression prediction from sequence by integrating long-range interactions. Nat. Methods 18, 1196–1203 (2021).

35. Sinai, S. & Kelsic, E. D. A primer on model-guided exploration of fitness landscapes for biological sequence design. arXiv [q-bio.QM] (2020).

36. Linder, J. & Seelig, G. Fast activation maximization for molecular sequence design. BMC Bioinformatics 22, 510 (2021).

37. Sinai, S., et al. AdaLead: A simple and robust adaptive greedy search algorithm for sequence design. arXiv *[cs.LG]* (2020).

38. Zrimec, J. et al. Controlling gene expression with deep generative design of regulatory DNA. Nat. Commun. 13, 5099 (2022).

39. Gupta, A. & Kundaje, A. Targeted optimization of regulatory DNA sequences with neural editing architectures. bioRxiv 714402 (2019) doi:10.1101/714402.

40. Killoran, N., Lee, L. J., Delong, A., Duvenaud, D. & Frey, B. J. Generating and designing DNA with deep generative models. arXiv [cs.LG] (2017).

41. Deverman, B. E., Ravina, B. M., Bankiewicz, K. S., Paul, S. M. & Sah, D. W. Y. Gene therapy for neurological disorders: progress and prospects. Nat. Rev. Drug Discov. 17, 767 (2018).

42. Mitchell, M. J. et al. Engineering precision nanoparticles for drug delivery. Nat. Rev. Drug Discov. 20, 101–124 (2020).

43. Tabebordbar, M. et al. Directed evolution of a family of AAV capsid variants enabling potent muscle-directed gene delivery across species. Cell 184, 4919–4938.e22 (2021).

44. Morales, L., Gambhir, Y., Bennett, J. & Stedman, H. H. Broader Implications of Progressive Liver Dysfunction and Lethal Sepsis in Two Boys following Systemic High-Dose AAV. Mol. Ther. 28, 1753–1755 (2020).

45. Hinderer, C. et al. Severe Toxicity in Nonhuman Primates and Piglets Following High-Dose Intravenous Administration of an Adeno-Associated Virus Vector Expressing Human SMN. Hum. Gene Ther. 29, 285–298 (2018).

46. Kelley, D. R., Snoek, J. & Rinn, J. L. Basset: learning the regulatory code of the accessible genome with deep convolutional neural networks. Genome Res. 26, 990–999 (2016).

47. Cazares, T. A. et al. maxATAC: Genome-scale transcription-factor binding prediction from ATAC-seq with deep neural networks. PLoS Comput. Biol. 19, e1010863 (2023).

48. Locatelli, F. et al. Lentiglobin Gene Therapy for Patients with Transfusion-Dependent β-Thalassemia (TDT): Results from the Phase 3 Northstar-2 and Northstar-3 Studies. Blood 132, 1025 (2018).

49. Locatelli, F. et al. Betibeglogene Autotemcel Gene Therapy for Non–β0/β0 Genotype β-Thalassemia. N. Engl. J. Med. 386, 415–427 (2022).

50. Wong, R. L. et al. Lentiviral gene therapy for X-linked chronic granulomatous disease recapitulates endogenous CYBB regulation and expression. Blood 141, 1007–1022 (2023).

51. Kohn, D. B. et al. Lentiviral gene therapy for X-linked chronic granulomatous disease. Nat. Med. 26, 200–206 (2020).

52. Mendell, J. R. et al. Single-Dose Gene-Replacement Therapy for Spinal Muscular Atrophy. N. Engl. J. Med. 377, 1713–1722 (2017).

53. Siders, W. M. et al. Cytotoxic T lymphocyte responses to transgene product, not adeno-associated viral capsid protein, limit transgene expression in mice. Hum. Gene Ther. 20, 11–20 (2009).

54. Tao, N. et al. Sequestration of adenoviral vector by Kupffer cells leads to a nonlinear dose response of transduction in liver. Mol. Ther. 3, 28–35 (2001).

55. Ganesan, L. P. et al. Rapid and efficient clearance of blood-borne virus by liver sinusoidal endothelium. PLoS Pathog. 7, e1002281 (2011).

56. Golovin, D. et al. Google Vizier: A Service for Black-Box Optimization. in Proceedings of the 23rd ACM SIGKDD International Conference on Knowledge Discovery and Data Mining 1487–1495 (Association for Computing Machinery, 2017).

57. Snoek, J., Larochelle, H. & Adams, R. P. Practical bayesian optimization of machine learning algorithms. Adv. Neural Inf. Process. Syst. 25, (2012).

58. Thurman, R. E. et al. The accessible chromatin landscape of the human genome. Nature 489, 75–82 (2012).

59. Zhang, J. et al. An integrative ENCODE resource for cancer genomics. Nat. Commun. 11, 3696 (2020).

60. Hardison, R. C. & Taylor, J. Genomic approaches towards finding cis-regulatory modules in animals. Nat. Rev. Genet. 13, 469–483 (2012).

61. Liu, Y. et al. Functional assessment of human enhancer activities using whole-genome STARR-sequencing. Genome Biol. 18, 219 (2017).

62. Luo, Y. et al. New developments on the Encyclopedia of DNA Elements (ENCODE) data portal. Nucleic Acids Res. 48, D882–D889 (2020).

63. Kagda, M. S. et al. Data navigation on the ENCODE portal. arXiv [q-bio.GN] (2023).

64. Hitz, B. C., et al. The ENCODE Uniform Analysis Pipelines. bioRxiv (2023) doi:10.1101/2023.04.04.535623.

65. van Laarhoven, P. J. M. & Aarts, E. H. L. Simulated annealing. in Simulated Annealing: Theory and Applications (eds. van Laarhoven, P. J. M. & Aarts, E. H. L.) 7–15 (Springer Netherlands, 1987).

66. Castro-Mondragon, J. A., et al. JASPAR 2022: the 9th release of the open-access database of transcription factor binding profiles. Nucleic Acids Res. 50, D165–D173 (2022).

67. Kulakovskiy, I. V. et al. HOCOMOCO: towards a complete collection of transcription factor binding models for human and mouse via large-scale ChIP-Seq analysis. Nucleic Acids Res. 46, D252–D259 (2018).

68. Sundararajan, M., Taly, A. & Yan, Q. Axiomatic Attribution for Deep Networks. in Proceedings of the 34th International Conference on Machine Learning (eds. Precup, D. & Teh, Y. W.) vol. 70 3319–3328 (PMLR, 06--11 Aug 2017).

69. Fulco, C. P. et al. Systematic mapping of functional enhancer–promoter connections with CRISPR interference. Science 354, 769–773 (2016).

70. Parviz, F. et al. Hepatocyte nuclear factor 4alpha controls the development of a hepatic epithelium and liver morphogenesis. Nat. Genet. 34, 292–296 (2003).

71. Harries, L. W., Brown, J. E. & Gloyn, A. L. Species-specific differences in the expression of the HNF1A, HNF1B and HNF4A genes. PLoS One 4, e7855 (2009).

72. El-Khairi, R. & Vallier, L. The role of hepatocyte nuclear factor 1β in disease and development. Diabetes Obes. Metab. 18 Suppl 1, 23–32 (2016).

73. Odom, D. T. et al. Core transcriptional regulatory circuitry in human hepatocytes. Mol. Syst. Biol. 2, 2006.0017 (2006).

74. Zweidler-Mckay, P. A., Grimes, H. L., Flubacher, M. M. & Tsichlis, P. N. Gfi-1 encodes a nuclear zinc finger protein that binds DNA and functions as a transcriptional repressor. Mol. Cell. Biol. 16, 4024–4034 (1996).

75. Huang, D.-Y., Kuo, Y.-Y. & Chang, Z.-F. GATA-1 mediates auto-regulation of Gfi-1B transcription in K562 cells. Nucleic Acids Res. 33, 5331–5342 (2005).

76. Beauchemin, H. & Möröy, T. Multifaceted Actions of GFI1 and GFI1B in Hematopoietic Stem Cell Self-Renewal and Lineage Commitment. Front. Genet. 11, 591099 (2020).

77. Agoston, Z. & Schulte, D. Meis2 competes with the Groucho co-repressor Tle4 for binding to Otx2 and specifies tectal fate without induction of a secondary midbrain-hindbrain boundary organizer. Development 136, 3311–3322 (2009).

78. Machon, O., Masek, J., Machonova, O., Krauss, S. & Kozmik, Z. Meis2 is essential for cranial and cardiac neural crest development. BMC Dev. Biol. 15, 40 (2015).

79. Zha, Y. et al. MEIS2 is essential for neuroblastoma cell survival and proliferation by transcriptional control of M-phase progression. Cell Death Dis. 5, e1417 (2014).

80. Lee, D. D. & Seung, H. S. Learning the parts of objects by non-negative matrix factorization. Nature 401, 788–791 (1999).

81. Birnbaum, R. Y. et al. Coding exons function as tissue-specific enhancers of nearby genes. Genome Res. 22, 1059–1068 (2012).

82. Kvon, E. Z. et al. Comprehensive In Vivo Interrogation Reveals Phenotypic Impact of Human Enhancer Variants. Cell 180, 1262–1271.e15 (2020).

83. Chatterjee, R. et al. Overlapping ETS and CRE Motifs ((G/C)CGGAAGTGACGTCA) preferentially bound by GABPα and CREB proteins. G3 2, 1243–1256 (2012).

84. Farley, E. K., Olson, K. M., Zhang, W., Rokhsar, D. S. & Levine, M. S. Syntax compensates for poor binding sites to encode tissue specificity of developmental enhancers. Proceedings of the National Academy of Sciences of the United States of America vol. 113 6508–6513 (2016).

85. Farley, E. K. et al. Suboptimization of developmental enhancers. Science 350, 325–328 (2015).

86. Cock, P. J. A. et al. Biopython: freely available Python tools for computational molecular biology and bioinformatics. Bioinformatics 25, 1422–1423 (2009).

87. Quinlan, A. R. & Hall, I. M. BEDTools: a flexible suite of utilities for comparing genomic features. Bioinformatics 26, 841–842 (2010).

88. Dale, R. K., Pedersen, B. S. & Quinlan, A. R. Pybedtools: a flexible Python library for manipulating genomic datasets and annotations. Bioinformatics 27, 3423–3424 (2011).

89. Heinz, S. et al. Simple combinations of lineage-determining transcription factors prime cis-regulatory elements required for macrophage and B cell identities. Mol. Cell 38, 576–589 (2010).

90. Bailey, T. L., Johnson, J., Grant, C. E. & Noble, W. S. The MEME Suite. Nucleic Acids Res. 43, W39–49 (2015).

91. Bailey, T. L. STREME: accurate and versatile sequence motif discovery. Bioinformatics 37, 2834–2840 (2021).

92. Gupta, S., Stamatoyannopoulos, J. A., Bailey, T. L. & Noble, W. S. Quantifying similarity between motifs. Genome Biol. 8, R24 (2007).

93. Santana-Garcia, W., et al. RSAT 2022: regulatory sequence analysis tools. Nucleic Acids Res. 50, W670–W676 (2022).

94. Grant, C. E., Bailey, T. L. & Noble, W. S. FIMO: scanning for occurrences of a given motif. Bioinformatics 27, 1017–1018 (2011).

95. Owen, A. B. & Perry, P. O. Bi-cross-validation of the SVD and the nonnegative matrix factorization. aoas 3, 564–594 (2009).

96. Kawakami, K. et al. A transposon-mediated gene trap approach identifies developmentally regulated genes in zebrafish. Dev. Cell 7, 133–144 (2004).

