## Supplementary Figures for "Machine-guided design of synthetic cell type-specific *cis*-regulatory elements"

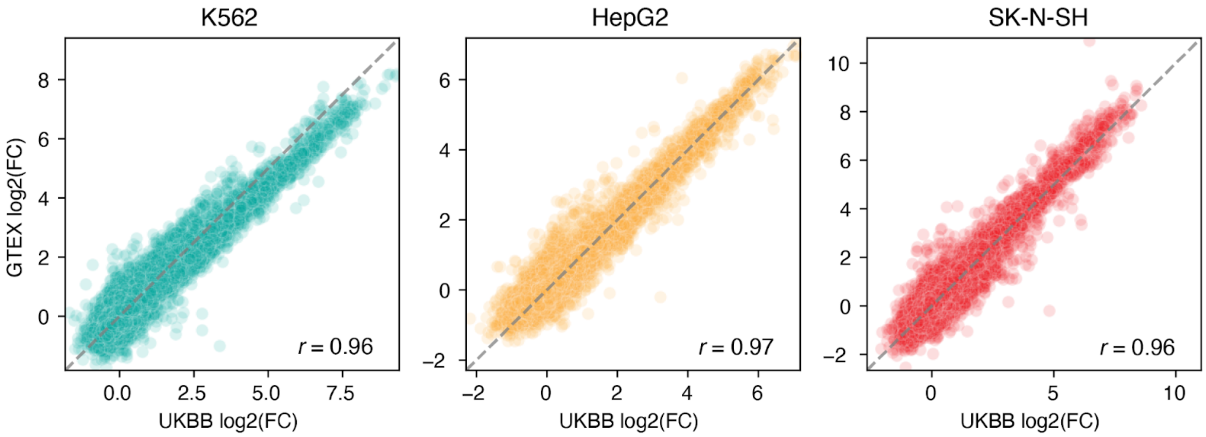


**Supplementary Figure 1. MPRA library reproducibility.** Scatter plots compare the $log_2$(Fold-Change) ($log_2$(FC)) of 20,303 sequences shared between the UKBB and GTEx MPRA libraries, two libraries experimentally conducted independently from each other at distinct points of time. The *x*-axis corresponds to the $log_2$(FC) as measured in UKBB, and the *y*-axis corresponds to the $log_2$(FC) as measured in GTEx. The Pearson's correlation coefficient is shown in the right bottom corner. Oligos with a replicate $log_2$(FC) standard error greater than 1 were omitted from the comparisons.

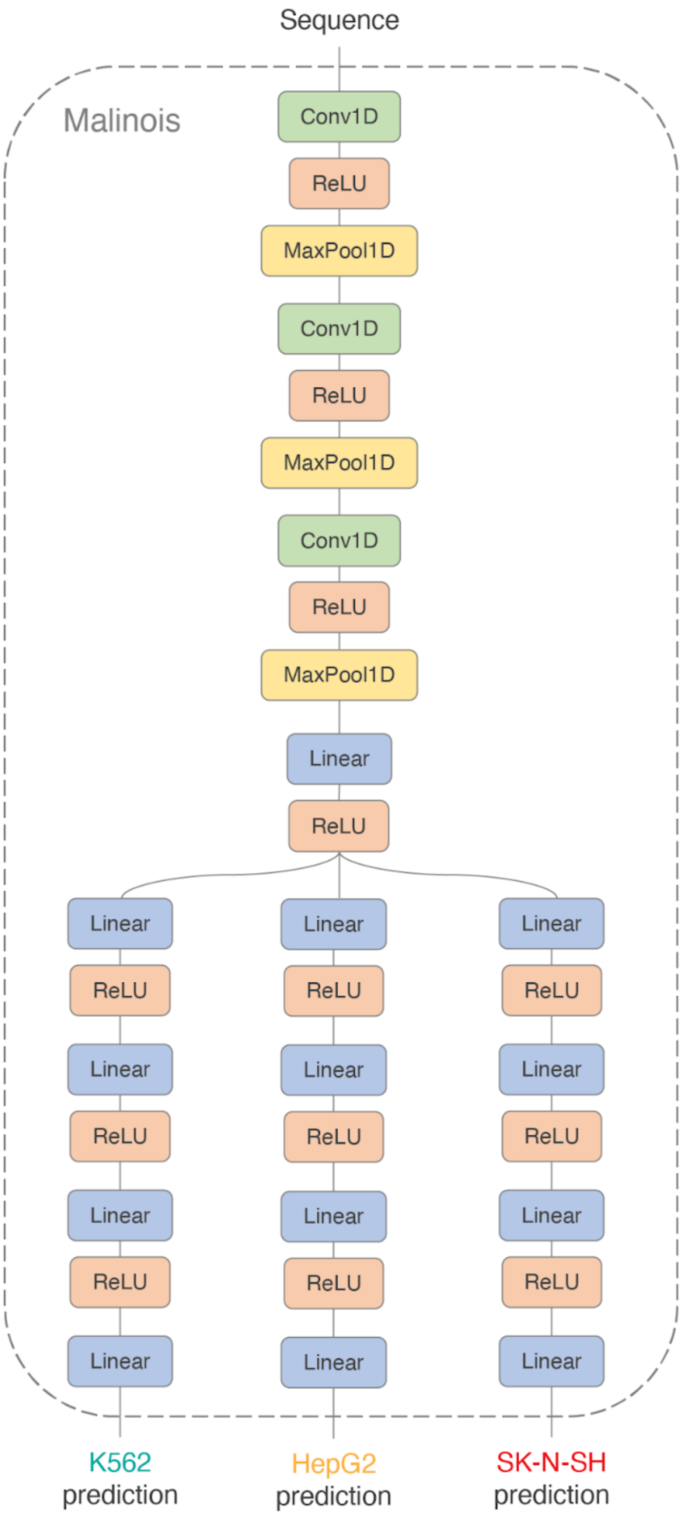


**Supplementary Figure 2. Model schematic.** Schematic of the Malinois model architecture. Malinois is composed of 3 convolutional layers, 1 shared linear layer, and 3 independent branches of 4 linear layers—1 branch for activity predictions in each cell type. All hidden layers are followed by rectified linear units while convolutional layers are also separated by pooling operations.

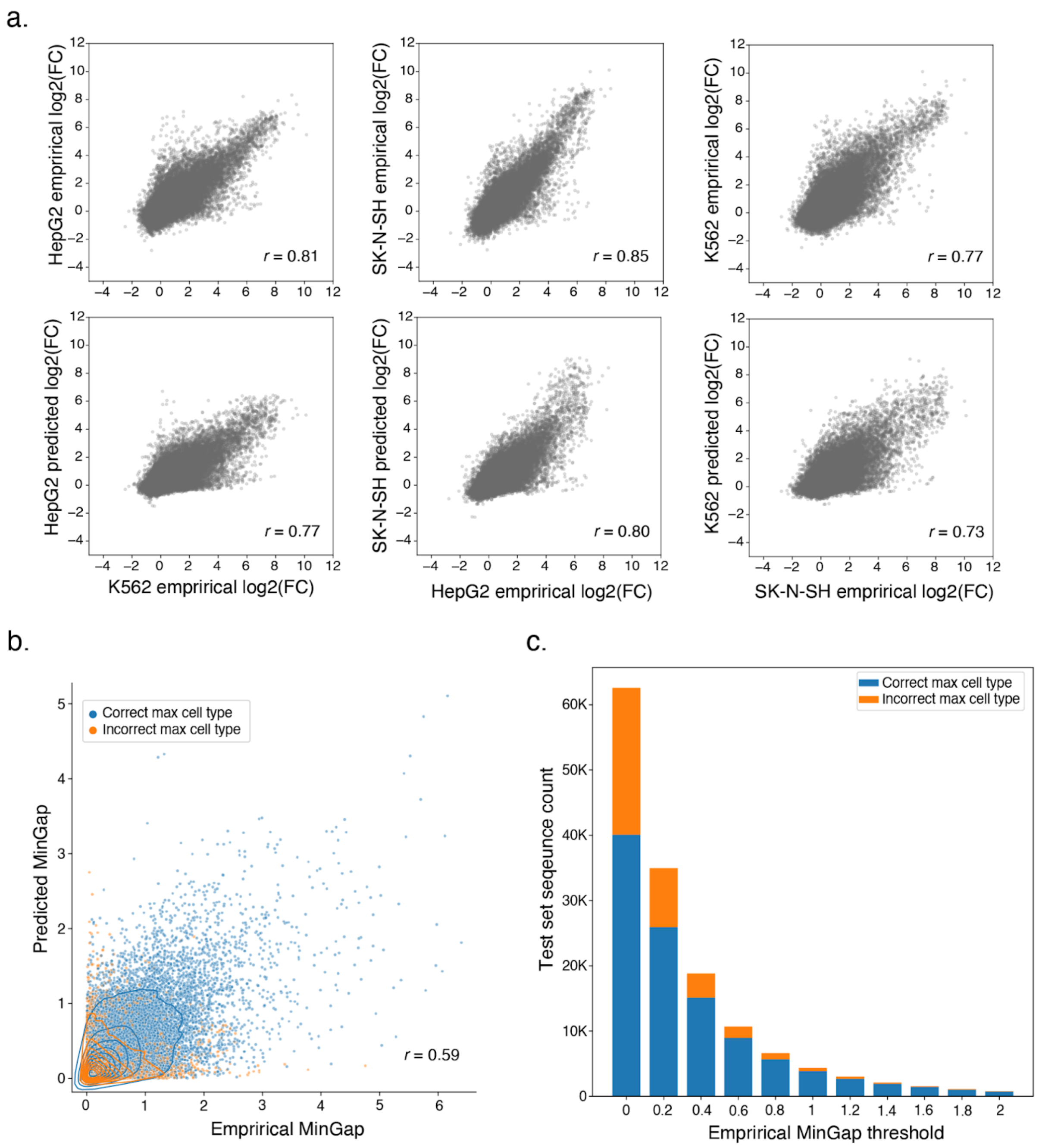


**Supplementary Figure 3. Cell type accuracy of model.** (**a**) Cross cell-type activity comparisons between empirical measurements and Malinois predictions organize and correlate similarly to empirical-to-empirical comparisons. Top scatter plots: empirical vs empirical cross-cell-type log2(FC). Bottom scatter plots: empirical vs predicted cross-cell-type log2(FC). Pearson correlation coefficients are shown in the left-bottom corner of each scatter plot. (**b**) Malinois can be used to identify highly active cell type-specific CREs. MinGap scores calculated using Malinois predictions correlate well with MPRA MinGap measurements for sequences in the held-out test set. Points are colored based on correct prediction of maximally active cell type by Malinois. (**c**) Malinois predictions of cell type associated with maximum CRE function are more accurate for sequences with high empirical specificity. Stacked bar plot displaying number of sequences in the test set falling into discrete bins based on an empirically measured MinGap threshold. Lower boundary of each bin is indicated on the *x*-axis and hue delineates sequences that are categorized correctly (blue) or incorrectly (orange).

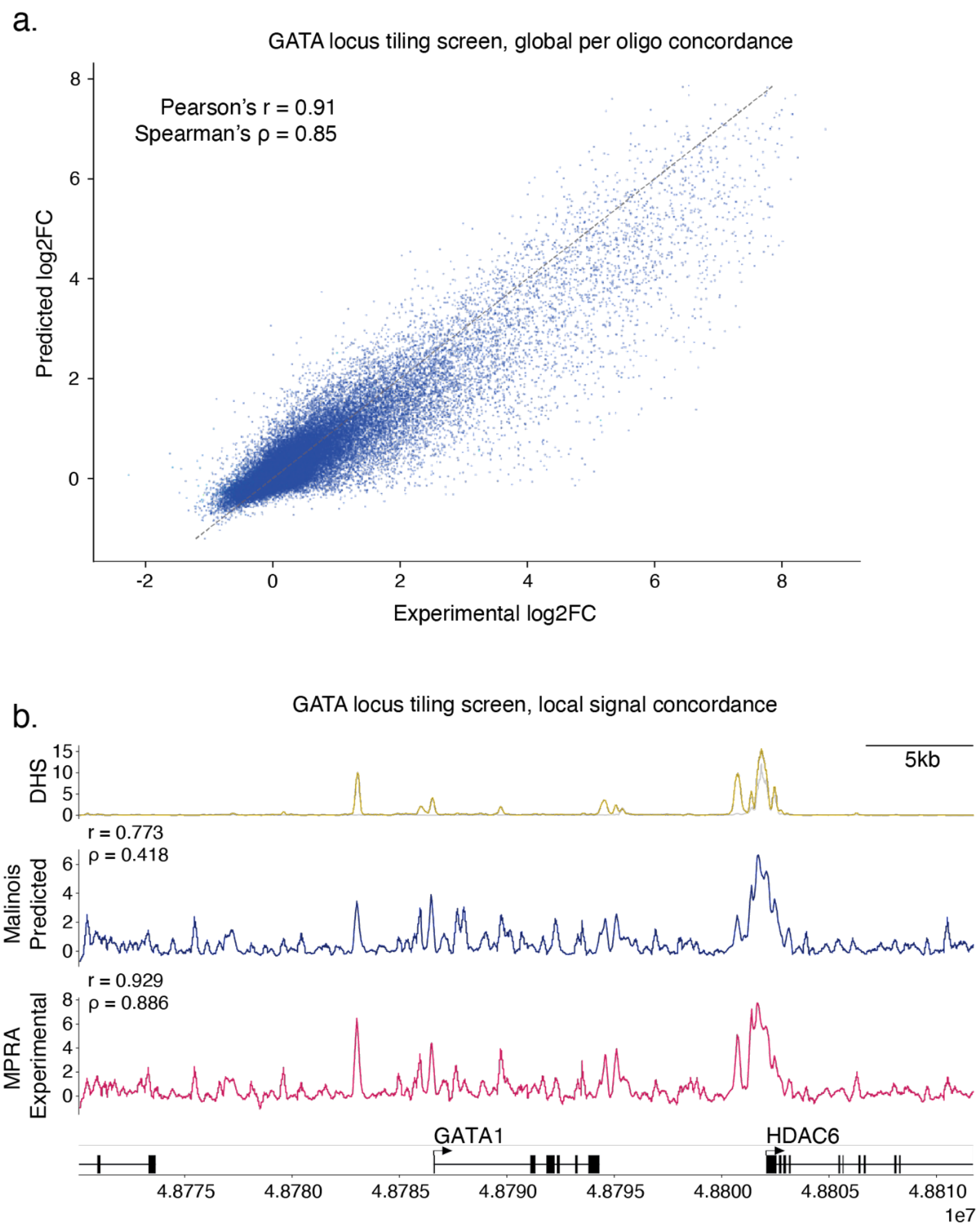


**Supplementary Figure 4. Correlation of Malinois predictions and empirical MPRA tiling data.** (**a**) Malinois predictions are highly correlated with empirical MPRA measurements of tiled sequences in the GATA locus (chrX:47,785,602:49,880,397)[5,48–50] in K562 with a Pearson's $r$ = 0.91. *X*-axis and *y*-axis correspond to empirical measurements and Malinois predictions, respectively for oligos in the library. Sequences with a replicate $\log_2$FC standard error greater than 1 in any cell type were omitted from the plots. (**b**) Malinois predictions projected onto the genome are correlated with empirical MPRA projections and DHS signal in regions with active CREs. Pearson's $r$ and Spearman's rho are calculated for the predicted track compared to either DHS (upper) or MPRA (lower).

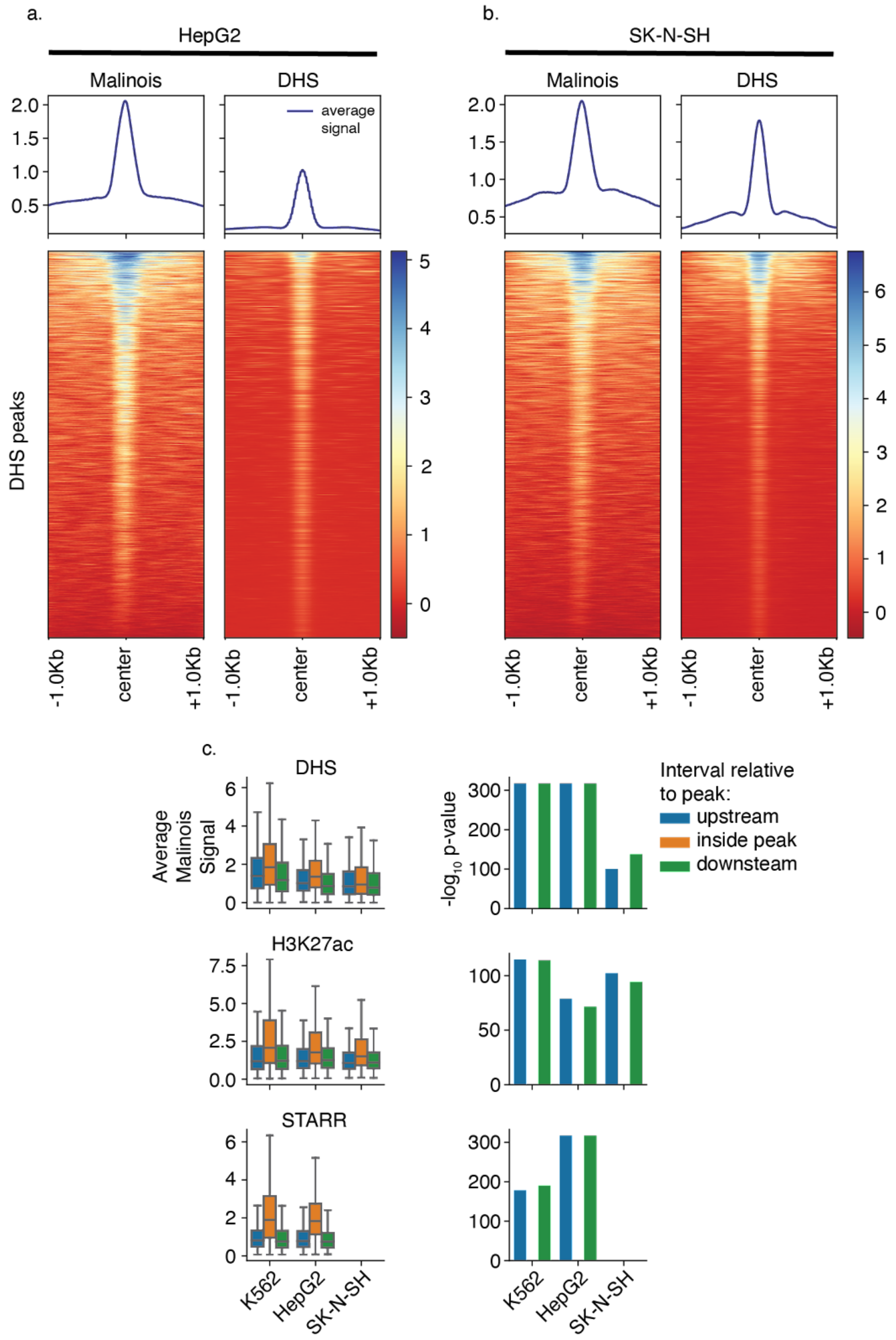


**Supplementary Figure 5. Malinois concordance with DHS/H3K27ac/STARR.** (**a**) Malinois genome-wide predictions correspond well with DHS signal in HepG2. Deeptools plots of Malinois genome-wide predictions and DHS signal centered at DHS peaks in HepG2 cell lines on chromosome 13. (**b**) DHS signal and Malinois genome-wide predictions are also similar in SK-N-SH. Similar Deeptools plots to **a** except using SK-N-SH derived data. (**c**) Malinois genome-wide predictions are significantly associated with candidate CRE mapping (DHS-seq, and H3K27ac ChIP-seq) and orthogonal signals of CRE functional characterization (STARR-seq). Boxplots (left) display average signal generated by Malinois genome-wide predictions within peaks annotated using alternative technologies (orange) compared to paired upstream (blue) and downstream (green) flanking regions. Associated bar plots (right) show *p*-values from paired *t*-tests comparing signals within peaks and outside of peaks. Coloring of bars indicates the flank used for comparison. Boxes demarcate the 25th, 50th, and 75th percentile values, while whiskers indicate the outermost point with 1.5 times the interquartile range from the edges of the boxes.

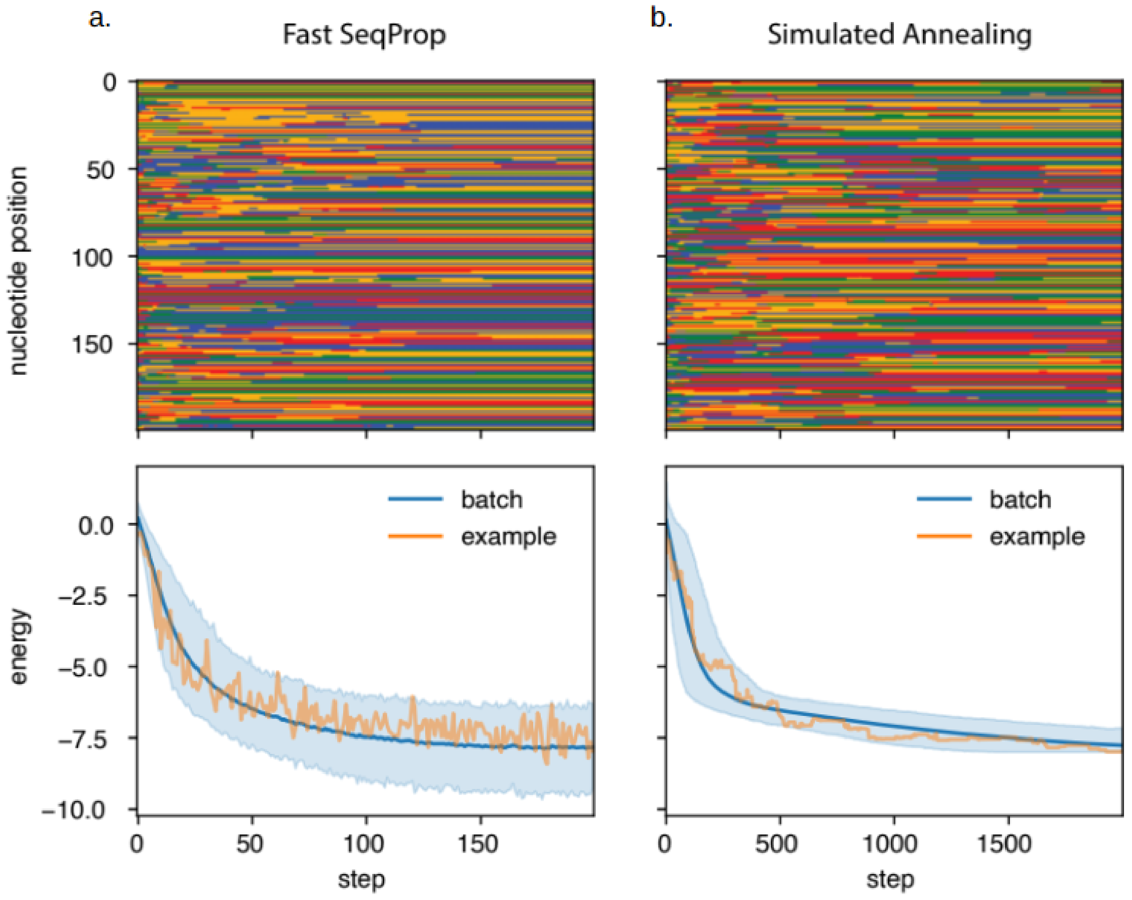


**Supplementary Figure 6. Example sequence generation trajectory.** (**a**) Fast SeqProp can generate sequences that are predicted to minimize an objective function. A trajectory was generated for 512 sequences using 200 update steps. Top: An example trajectory of a single sequence in the trajectory. Color represents nucleotide identity along the sequence after each update during the algorithm (A: Green, C: Blue, G: Yellow, T: Red). Bottom: The predicted objective value of sequences at each step of Fast SeqProp. The mean is indicated by the line and bounds of the 95 percentile data range are shaded light blue. The example displayed above is indicated by the orange line. (**b**) Same as **a**, but generated using 2000 steps of simulated annealing.

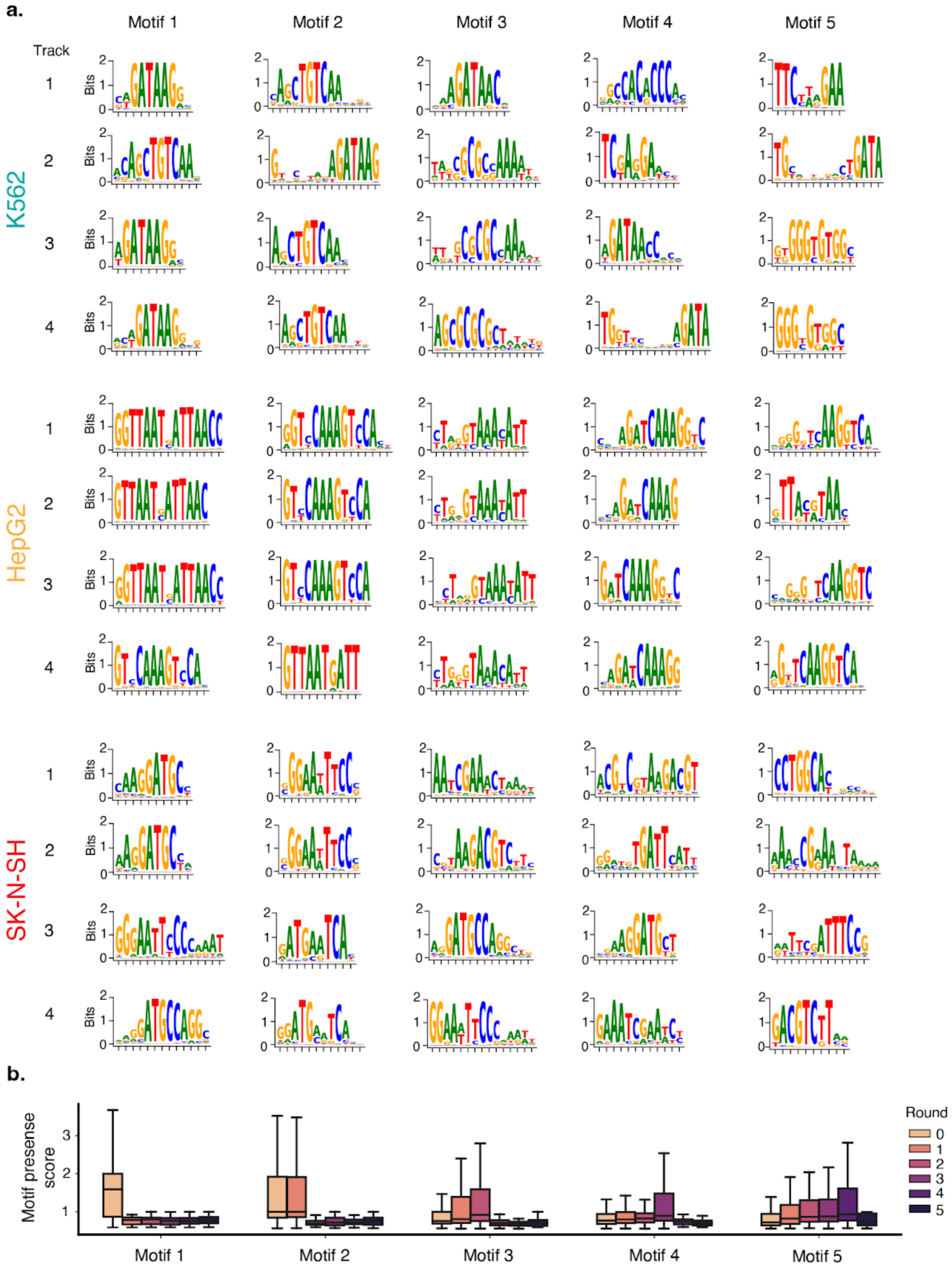


**Supplementary Figure 7. Motif match scores during penalization.** (**a**) Motifs can be depleted from Fast SeqProp-generated sequences using motif penalization. Motif numbers on the *x*-axis correspond to the first round in which their matches are penalized during Fast SeqProp, as they were the top match from the previous round. For each target cell type, four independent tracks of penalization were carried out (**Methods**) to account for potential enrichment effects of the random initialization when generating sequences. (**b**) Underrepresented motifs are progressively enriched as preferred alternatives are depleted. Box plots capture distribution of motif matches across sequences produced in each round of penalized generation. Motif numbers on the *x*-axis correspond to the first round in which their matches are penalized during Fast SeqProp. Motifs are specifically depleted in rounds where they are introduced into the penalty calculation, but can gradually rise during preceding rounds. In the *y*-axis, the motif-presence score of each motif is calculated by summing all the motif-match scores that pass a score threshold in a sequence, and dividing the sum by the score of the motif consensus sequence. Boxes demarcate the 25th, 50th, and 75th percentile values, while whiskers indicate the outermost point with 1.5 times the interquartile range from the edges of the boxes.

a.  b.

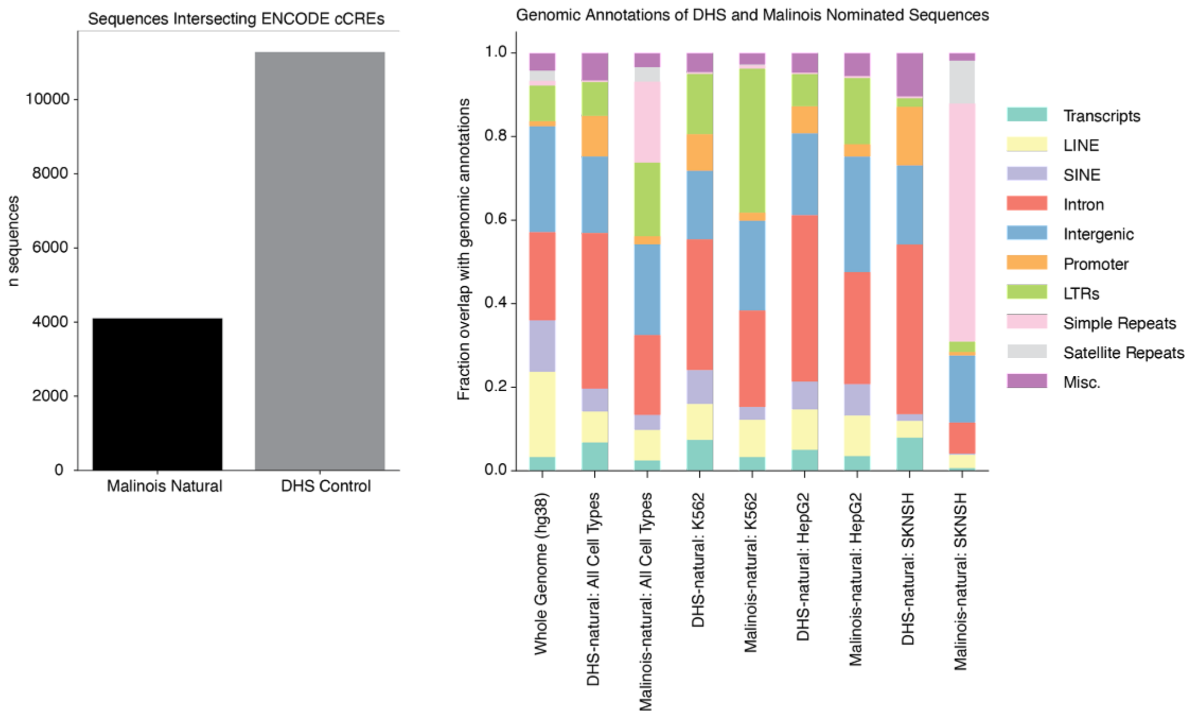


c.

| Malinois-natural \ DHS-natural | K562 | HepG2 | SK-N-SH |
|---|---|---|---|
| K562 | 154 | 1 | 2 |
| HepG2 | 0 | 162 | 1 |
| SK-N-SH | 0 | 2 | 4 |

**Supplementary Figure 8. Annotation of naturally occurring sequences.** (**a**) Sequences nominated by DHS accessibility (DHS-natural) and by Malinois (Malinois-natural) were intersected with ENCODE cCREs (promoter-like sequences, proximal enhancer-like sequences, distal enhancer-like sequences, and CTCF-only) to determine overlap with existing putative regulatory elements. 94% of DHS-natural sequences intersect a cCRE while only 34.2% of Malinois-natural sequences intersect a cCRE suggesting that Malinois may exploit sequences features not captured by typical cCRE measures to select a sequence that drives cell type-specific activity. (**b**) To explore additional genomic features that may overlap DHS-natural and Malinois-natural sequences were annotated using annotatePeaks.pl from the HOMER suite. Annotations were generated for the whole genome (hg38), the DHS-natural and Malinois-natural libraries as a whole, as well as DHS-natural and Malinois-natural by individual cell type. DHS-natural and Malinois-natural largely resemble the distribution of annotations genome-wide barring an overrepresentation of simple repeats in Malinois-natural sequences driven by SK-N-SH sequences. Despite this, selected sequences seem to be a representative sample of genomic features. (**c**) DHS-natural and Malinois-natural sequences were intersected to determine overlap between naturally occurring sequences. Notably overlap was minimal between selection methods (0.10%-4.1%) depending on cell type.

a.

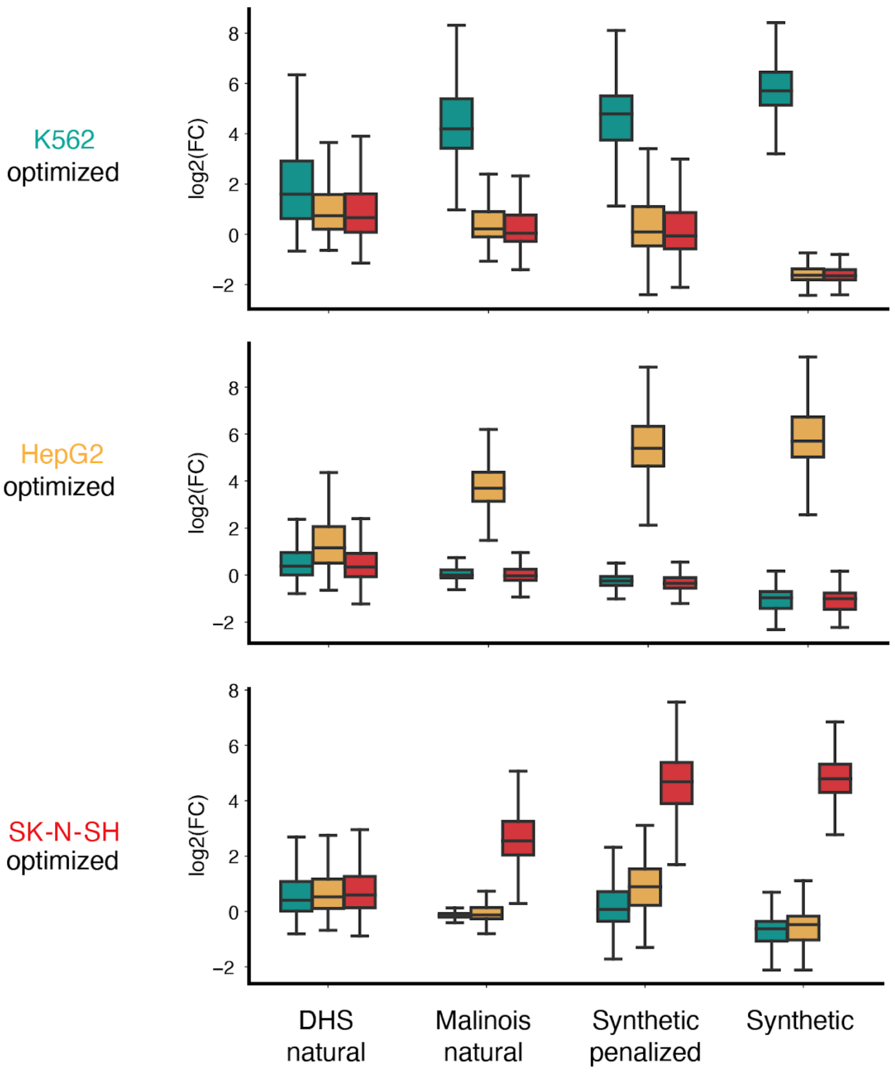


**Supplementary Figure 9 - Predicted library activity.** (**a**) Distribution of projected activity in K562 (teal), HepG2 (gold), and SK-N-SH (red) for candidate CREs predicted to drive K562-specific transcription. Boxes demarcate the 25th, 50th, and 75th percentile values, while whiskers indicate the outermost point with 1.5 times the interquartile range from the edges of the boxes. (**b**) Same as **a,** but for candidate CREs predicted to drive HepG2-specific transcription. (**c**) Same as **a** and **b,** but for candidate CREs predicted to drive SK-N-SH-specific transcription.

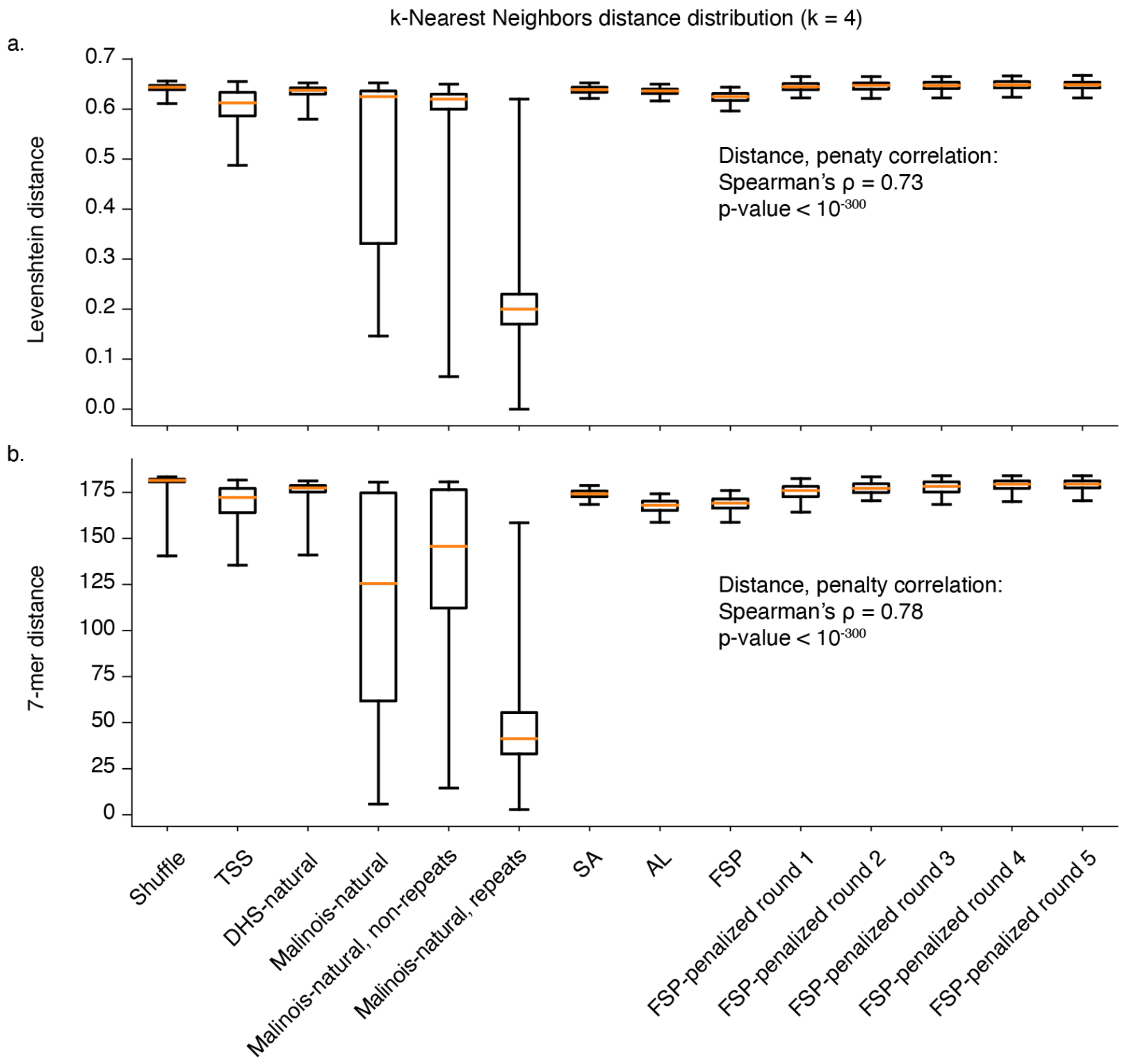


**Supplementary Figure 10. K-mer and Hamming distance.** (**a**) Algorithms for model-guided sequence designs produce diverse, non-degenerate candidate CREs. Box plot displays the distribution of average Levenshtein distance to 4 nearest neighbors for sequences in categories indicated on the *x*-axis. As a control, we randomly selected 4000 shuffled sequences from the candidate CRE library and 19381 promoter sequences extracted from RefGene by taking the 200 nucleotides upstream of (strand aware) TSS annotations for mRNAs. Malinois-natural results are plotted on aggregate, only using non-repeat element matched sequences, and repeat element matched sequences. Spearman's correlation coefficient was calculated between penalization round number (starting at zero) and average Hamming distances to 4 nearest neighbors. Boxes demarcate the 25th, 50th, and 75th percentile values, while whiskers indicate the 1st and 99th percentile values. (**b**) Algorithms for model-guided sequence designs produce sequences with diverse, non-redundant 7-mer usage. Plot is the same as **a** except it displays average L1 distance of 7-mer content between sequences and 4 nearest neighbors, divided by 2. Boxes demarcate the 25th, 50th, and 75th percentile values, while whiskers indicate the 1st and 99th percentile values.

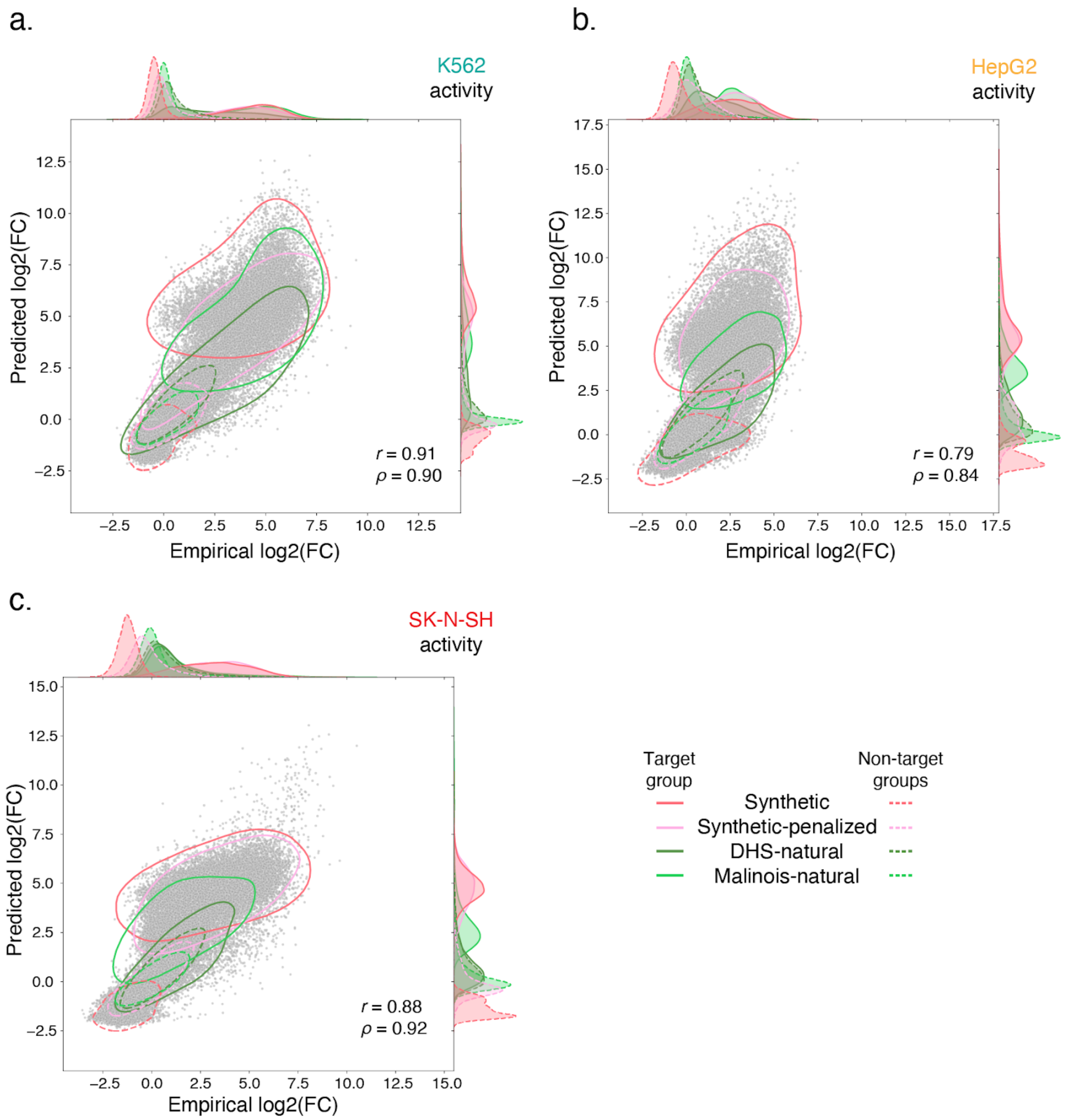


**Supplementary Figure 11. Library prediction validation plots.** (**a**) Prospective Malinois predictions of candidate cell type-specific CRE activity is correlated with experimental measurements across all three tested cell types. The scatter plot corresponds to predictions and measurements made in K562. Solid contour lines demarcate 95% density of points corresponding to candidate CRE expected to drive expression in K562. Dotted contour lines indicate 95% density of CREs expected to drive specific expression in one of the other two cell types. Color indicates sequence selection or generation method. One-dimensional density estimates along axes share the same line style and color associations. Sequences with a replicate $\log_2$FC standard error greater than 1 in any cell type were omitted from the plots. (**b**) Same as **a**, but in HepG2. (**c**) Same as **a**, but in SK-N-SH.

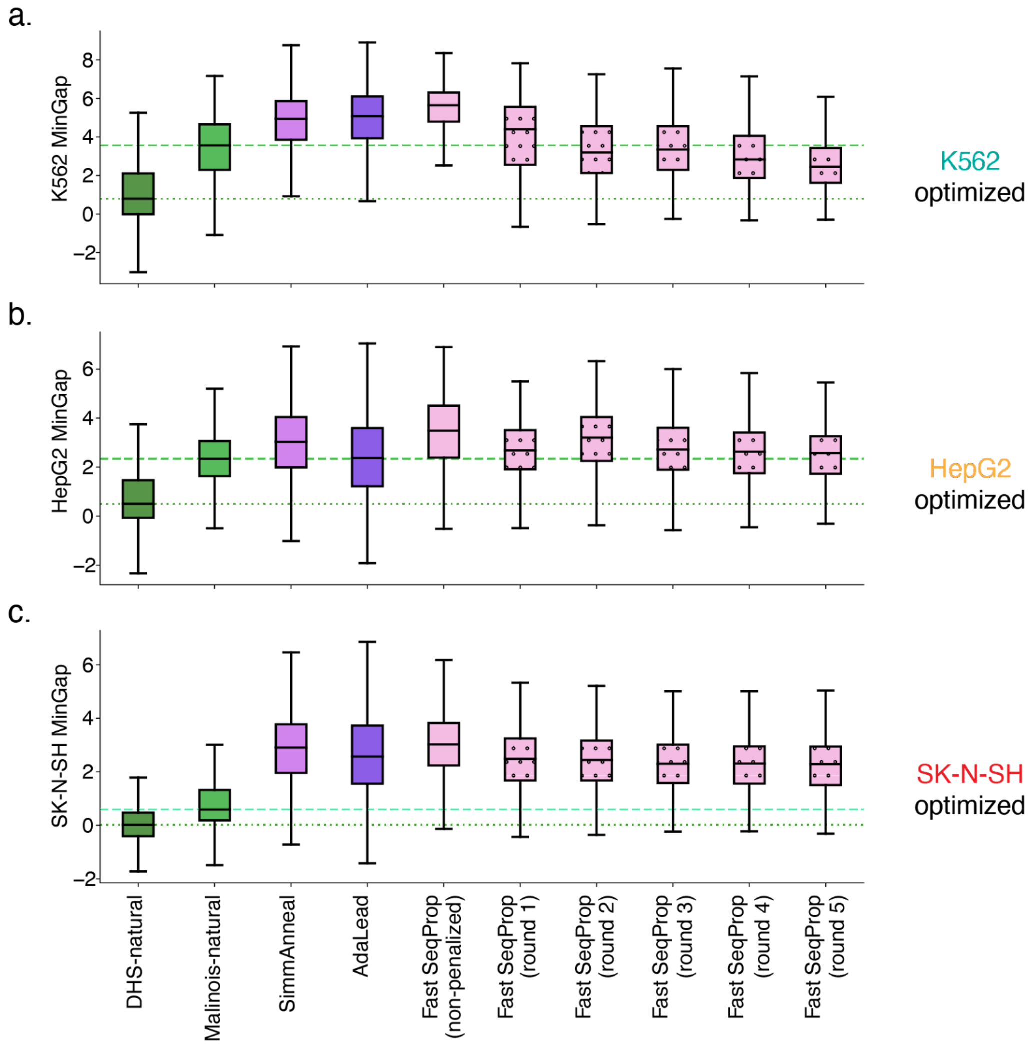


**Supplementary Figure 12. MinGap boxplots.** (**a**) Malinois improves identification of CREs with K562-specific activity and synthetic sequence generation enables creation of CREs with enhanced functions. Distribution of MPRA-measured K562-specific activity in various candidate CRE groups. Green and aquamarine lines indicate median MinGap of DHS-natural and Malinois-natural candidates respectively. Sequences with a replicate $\log_2$FC standard error greater than 1 in any cell type were omitted from the plots. Boxes demarcate the 25th, 50th, and 75th percentile values, while whiskers indicate the outermost point with 1.5 times the interquartile range from the edges of the boxes. (**b**) Same as a. except quantification of candidate sequences targeting HepG2. (**c**) Same as a. except quantification of candidate sequences targeting SK-N-SH.

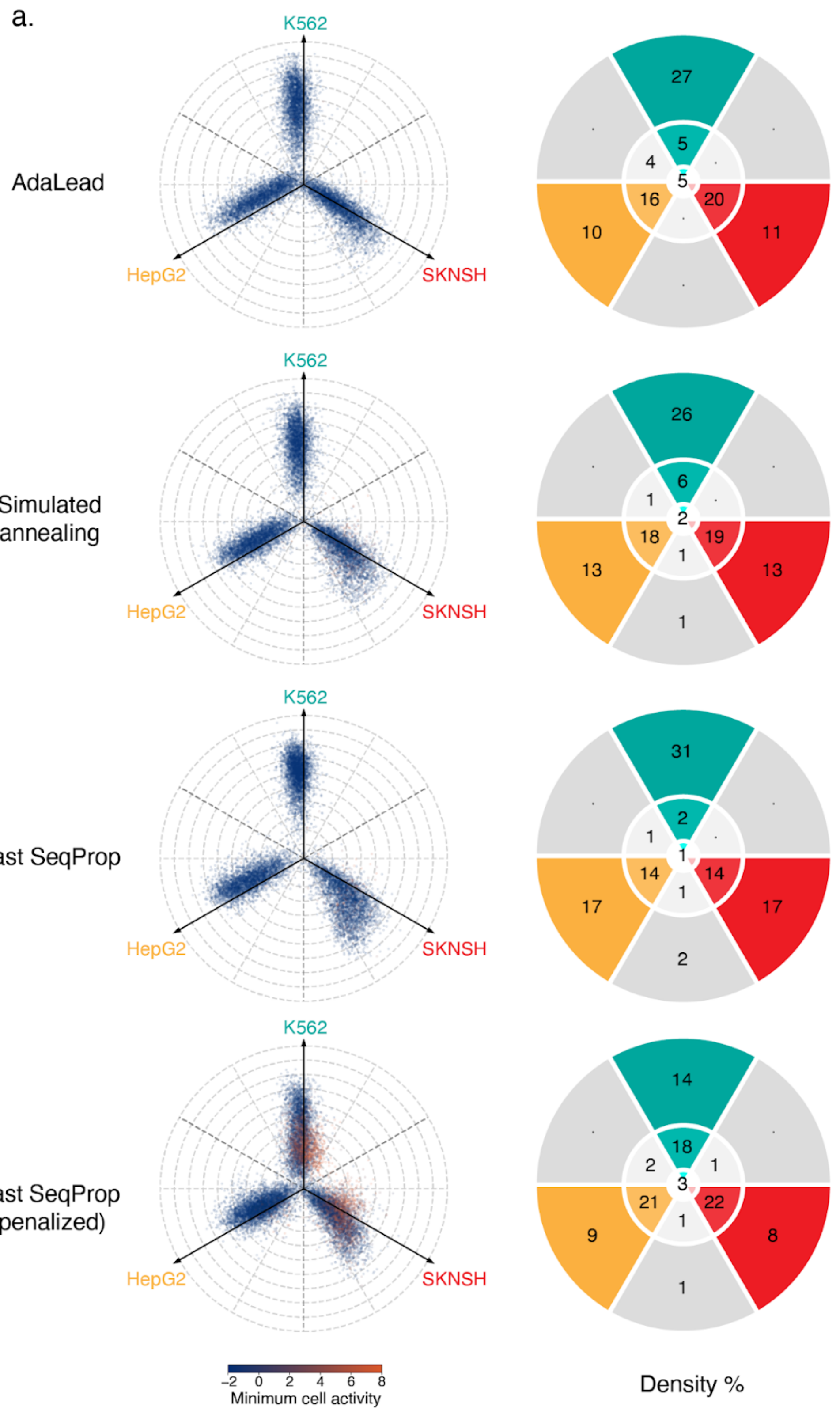


**Supplementary Figure 13. Complete propeller plots.** (**a**) Propeller plots of refined synthetic subsets of the library (see **Figure 2e** legend for description of coordinate system).

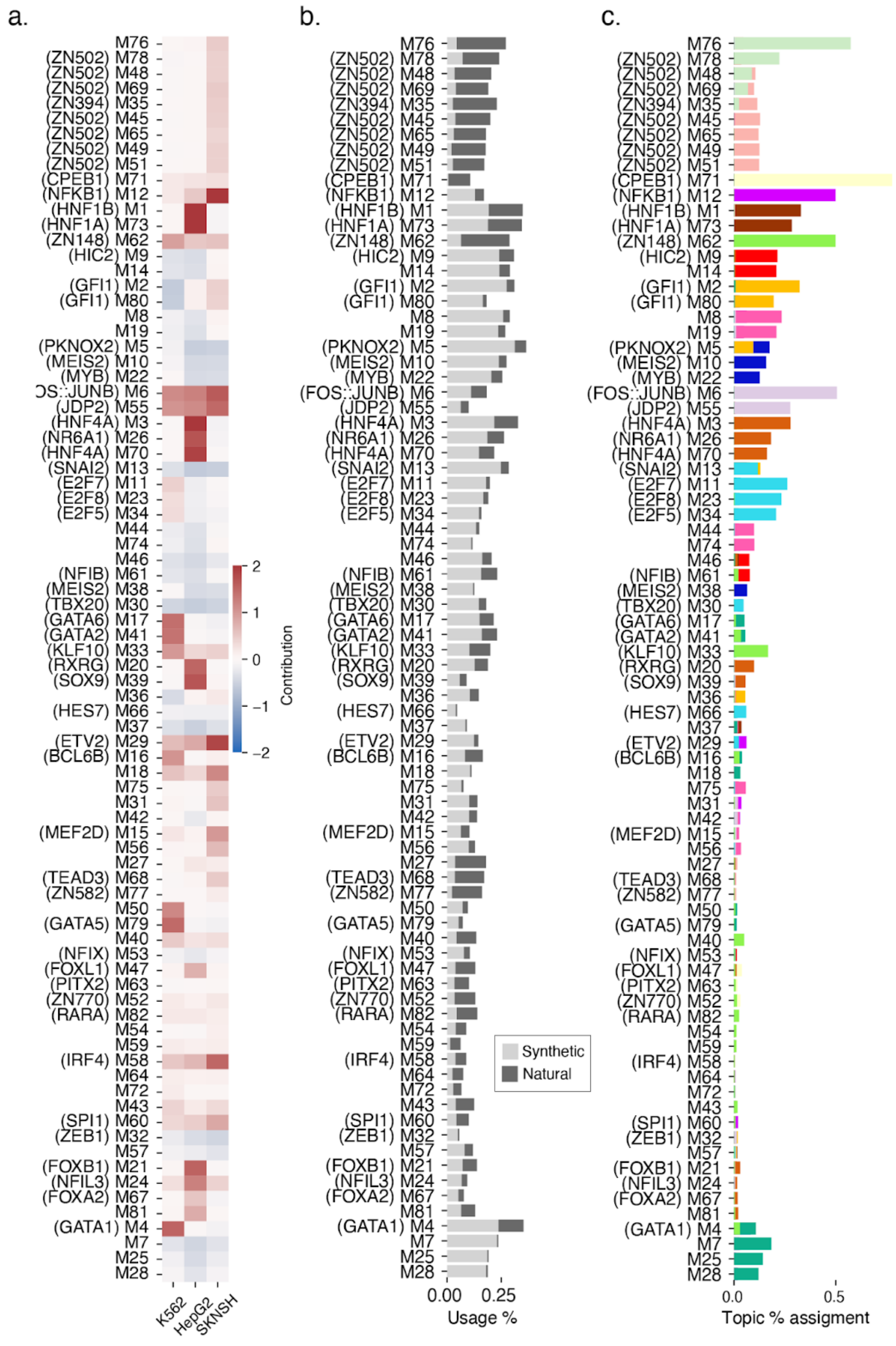


**Supplementary Figure 14. Motif associations to topics (bar-plots) with motif function summary.** (**a**) The 82 enriched motifs discovered in our collection of natural and synthetic cell type-specific CREs have diverse predicted patterns of function across K562, HepG2, and SK-N-SH. Heatmap quantifies average contribution scores across all motif matches in each cell type (columns, left to right: K562, HepG2, and SK-N-SH). Motifs are ordered along the *y*-axis based program associations in panel c. (**b**) Motif representation in the library is expressed as a fraction of sequences containing the motif. Coloring indicates if matched sequences are natural (dark gray) or synthetic (light gray). (**c**) Coefficients from the NMF feature matrix quantify strength of motif association with various higher order functional programs.

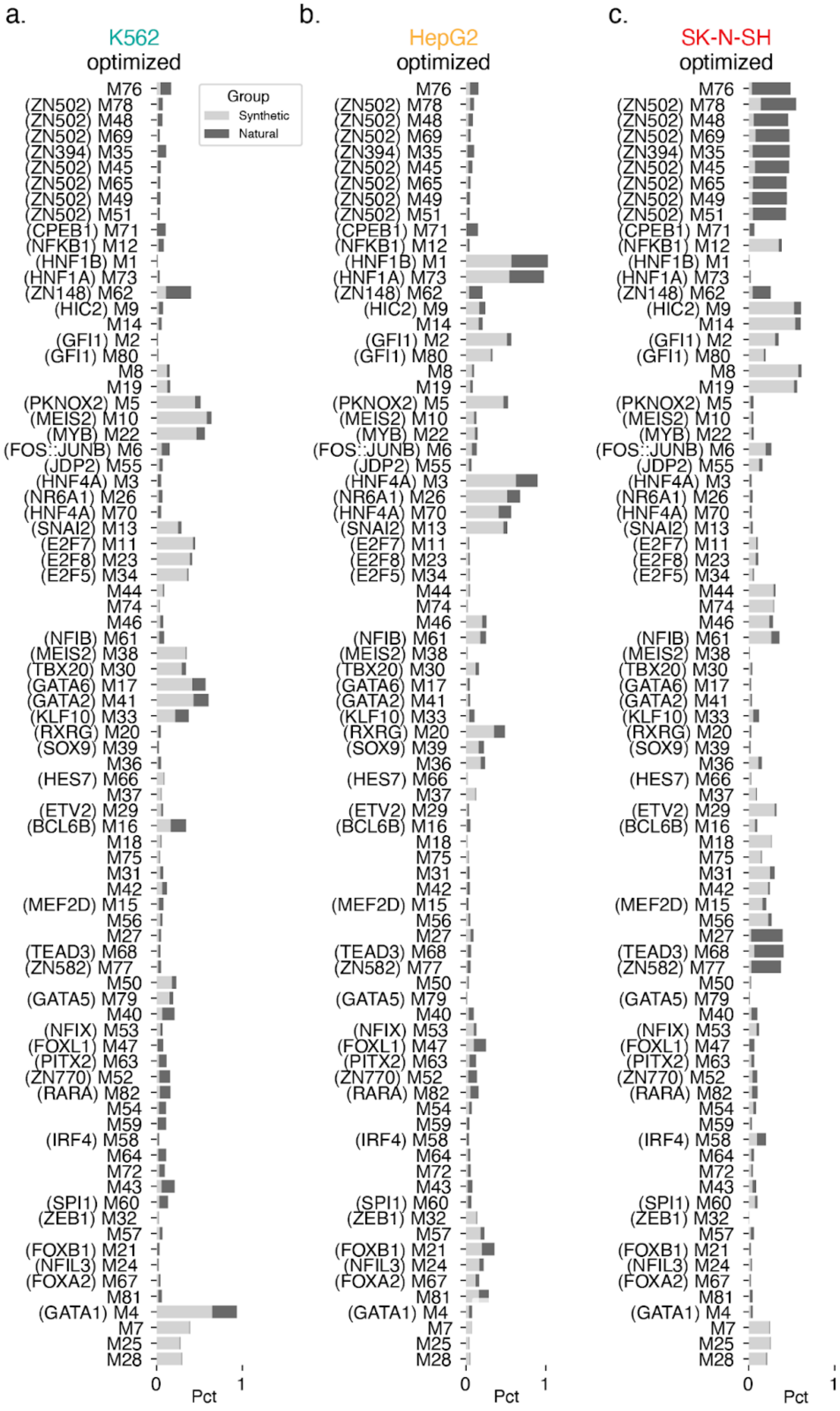


**Supplementary Figure 15. Motif enrichment by cell type target.** (**a**) Motif representation in K562-optimized sequences only. Bar width indicates the fraction of natural (dark gray) or synthetic (light gray) K562-optimized sequences containing the motif. (**b**) Same as **a**, but in HepG2-optimized. (**c**) Same as **a**, but in SK-N-SH-optimized.

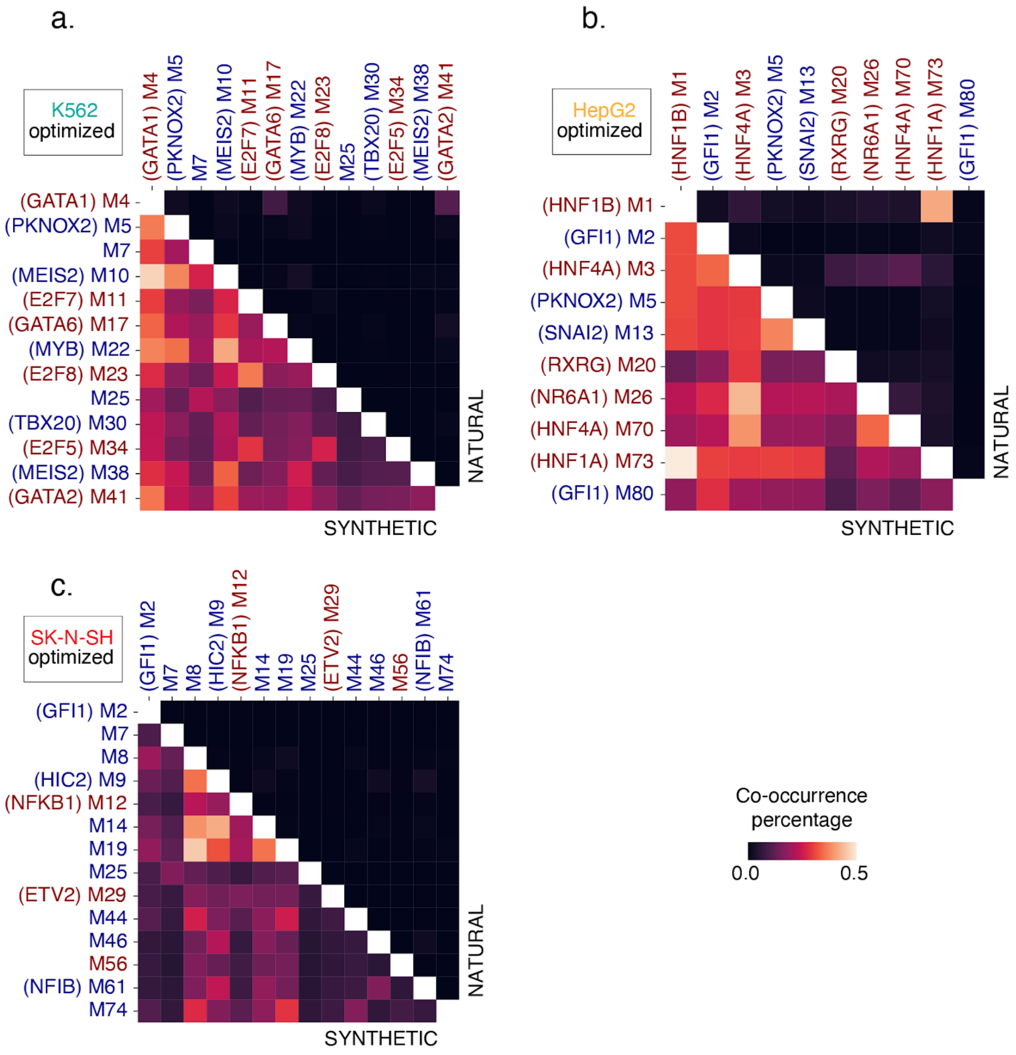


**Supplementary Figure 16. Uncorrected motif co-occurrence percentages.** (**a**) Motif co-occurrence representation in K562-optimized sequences only. Color indicates the fraction of natural (upper triangle) or synthetic (lower triangle) K562-optimized sequences containing a motif pair. (**b**) Same as **a**, but in HepG2-optimized. (**c**) Same as **a**, but in SK-N-SH-optimized.

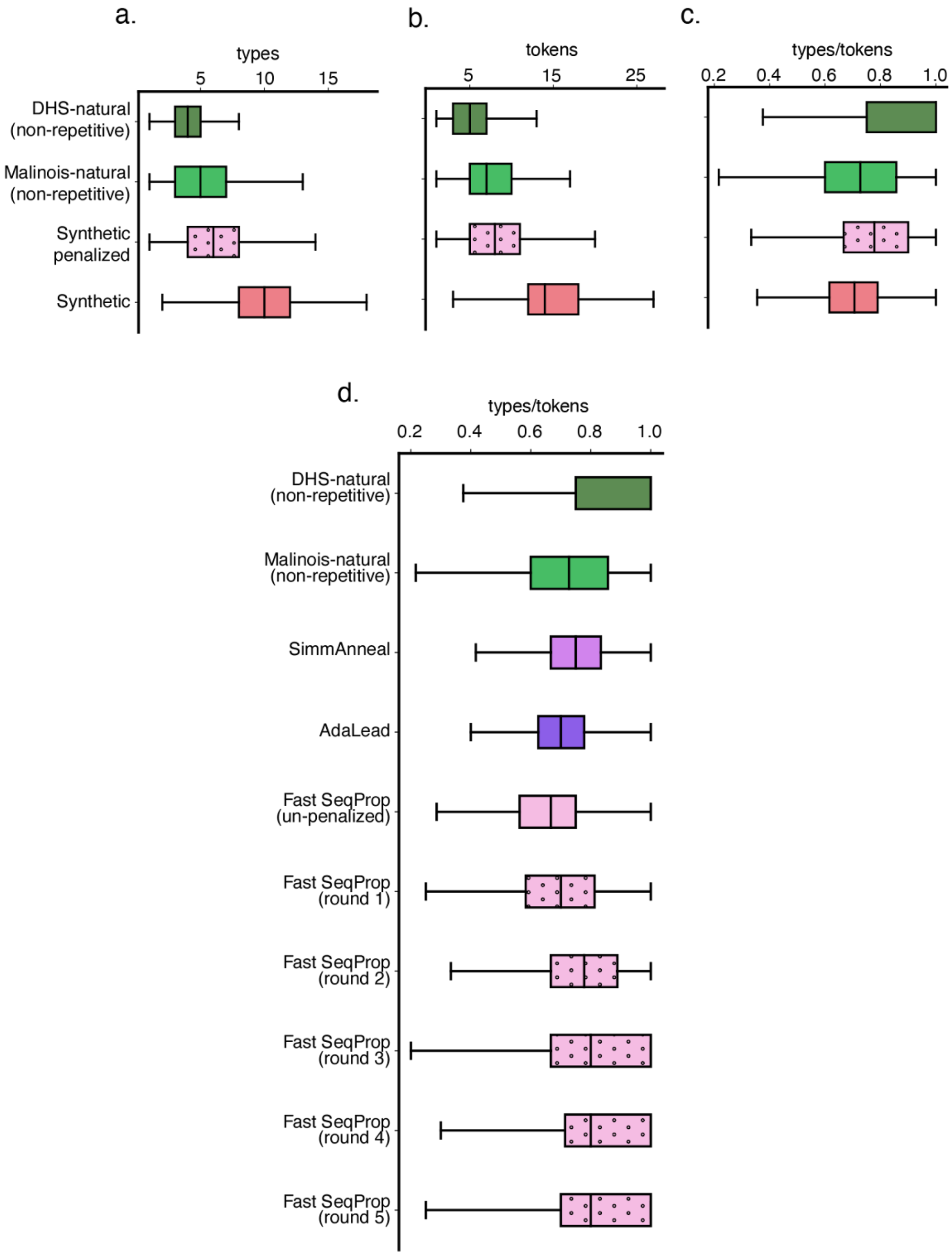


**Supplementary Figure 17. Type:token.** (**a**) Individual synthetic sequences are composed of more unique enriched sequence motifs than natural sequences. Distribution of unique motifs (types) in each sequence, binned by CRE proposal method. Boxes demarcate the 25th, 50th, and 75th percentile values, while whiskers indicate the outermost point with 1.5 times the interquartile range from the edges of the boxes. (**b**) Synthetic sequences contain more instances of enriched motifs than natural sequences. Distribution of total motif instances (tokens) in each sequence, binned by CRE proposal method. Boxes demarcate the 25th, 50th, and 75th percentile values, while whiskers indicate the outermost point with 1.5 times the interquartile range from the edges of the boxes. (**c**) Malinois-based CRE proposals contain more redundant motif matches compared to DHS-based CRE nominations. Distribution of type:token in each sequence, binned by CRE proposal method. Boxes demarcate the 25th, 50th, and 75th percentile values, while whiskers indicate the outermost point with 1.5 times the interquartile range from the edges of the boxes. (**d**) Motif penalization reduces motif redundancy in synthetic CREs. Boxplots are similar to c. except synthetic elements are broken up into more granular bins. Type:token ratio is significantly correlated with increasing penalization (Spearman's $\rho$=0.40, $p<10^{-300}$). Boxes demarcate the 25th, 50th, and 75th percentile values, while whiskers indicate the outermost point with 1.5 times the interquartile range from the edges of the boxes.

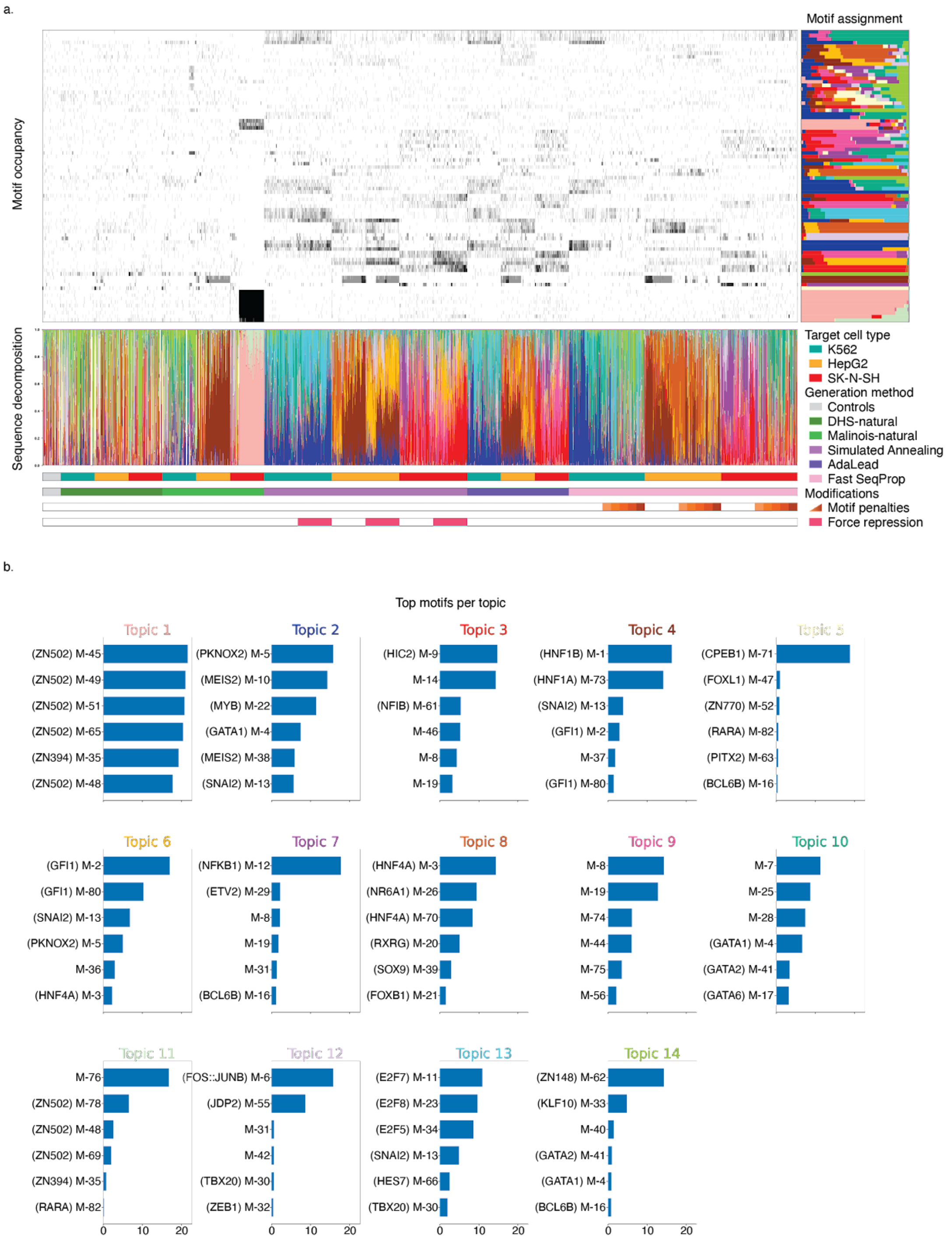


**Supplementary Figure 18. Full NMF structure plot and top-motif set per program.** (**a**) NMF decomposes sequence libraries and aggregates motifs into 14 distinct functional programs. Various CRE proposal methods favor distinct patterns of program usage. Top-left, grayscale heatmap: Motifs (*y*-axis) are identified in each sequence (*x*-axis). Shading indicates the number of motif matches in a sequence, capped at 5 matches. Top-right horizontal bar plot: Frequency of program association for each motif extracted from NMF feature matrix, unit normalized. *Y*-axis is shared with top-left and ordering was set by clustering motifs using the feature matrix. Topic coloring is consistent with **Figure 3d**. Bottom, vertical bar plot: Program decomposition of individual sequences, unit normalized. Bottom, colored stips: Demarcation of CRE metadata (i.e., predicted target cell type, generation method, objective function modification) with color corresponding to legend on the right and side. CREs are clustered within these subsets based on program content. (**b**) Raw values from the NMF feature matrix for the top 6 motifs associated with each topic. Coloring of topic subtitles is consistent with **Figure 3d**.

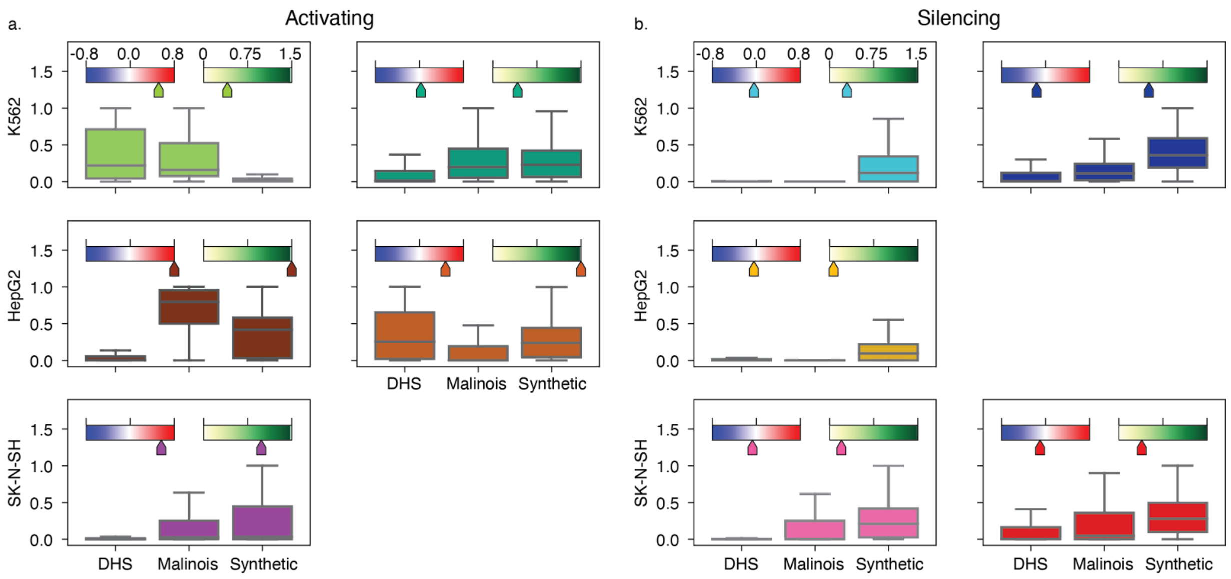


**Supplementary Figure 19. Activating, repressing, and ubiquitous program content and usage.** (**a**) Natural sequences that target K562- and HepG2-specific transcription favor activating programs. Program content distributions for 5 cell type specifying programs, plotted for the subset of sequences expected to target the cell type relevant to the program. These 5 programs display an average on-target activating effect. Blue-red slider indicates contribution scores at mapped motifs, weighted by relative program weight, and averaged over sequences and cell types to summarize on-target activating or off-target repressing activity of each program. White-green slider indicates overall degree of cell specification by program calculated from contribution scores at mapped motifs, weighted by relative program weight, and averaged over sequences followed by MinGap across cell types. Malinois-natural sequences overlapping with repetitive elements have been removed from this and following analysis of program content. Boxes demarcate the 25th, 50th, and 75th percentile values, while whiskers indicate the outermost point with 1.5 times the interquartile range from the edges of the boxes. (**b**) Repressive programs are preferred over activating to drive SK-N-SH-specific transcription for all groups of CREs. Similar program content plots to a. except showing programs with an average repressive effect in off-target cell types. Boxes demarcate the 25th, 50th, and 75th percentile values, while whiskers indicate the outermost point with 1.5 times the interquartile range from the edges of the boxes.

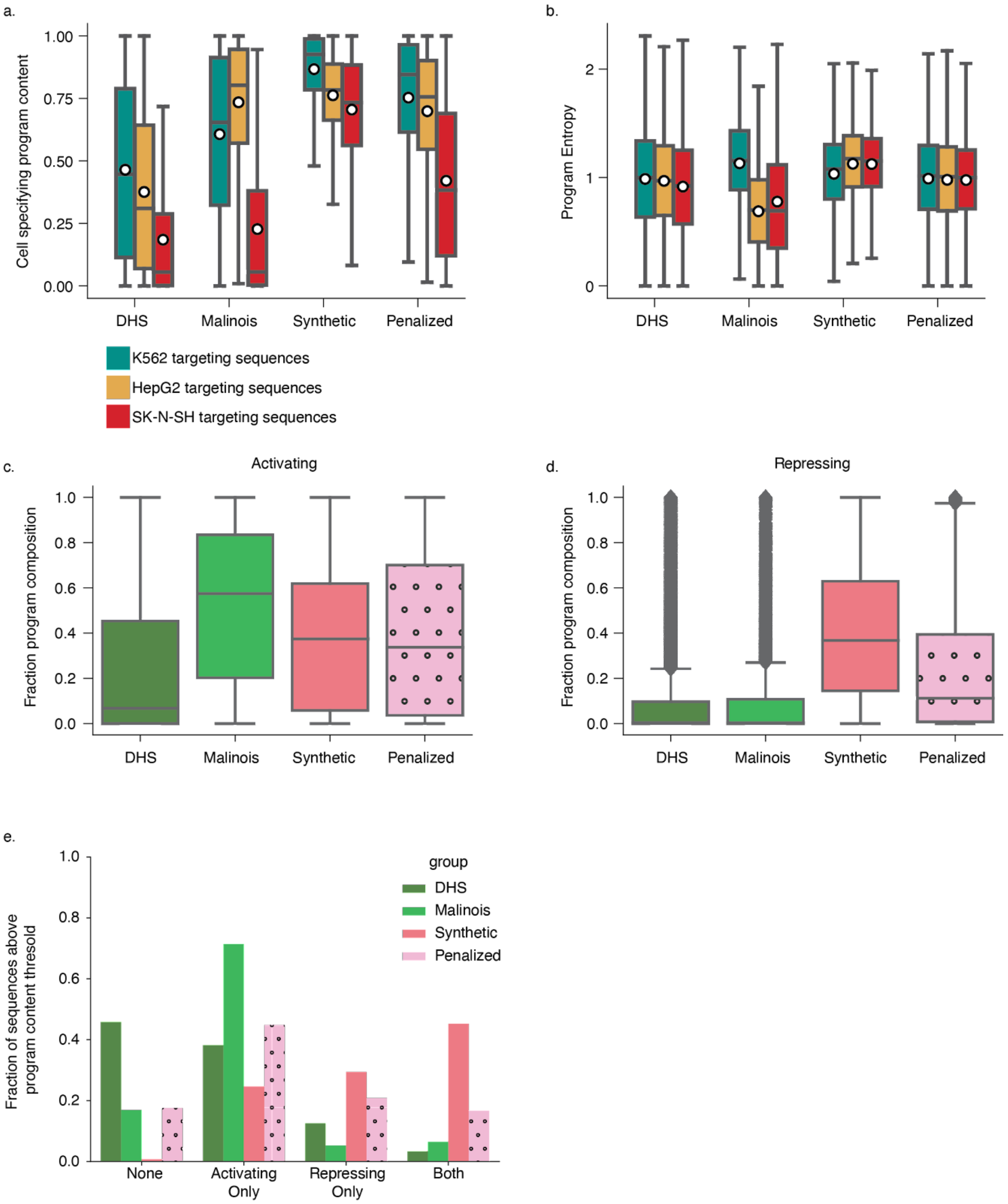


**Supplementary Figure 20. Overall program usage.** (**a**) Synthetic sequences utilize a greater density of cell type-specifying programs compared to natural sequences. Distribution of cell type-specifying program usage as a fraction of total program content in different subgroups of CREs. Welch's *t*-test $p<10^{-10}$ for all cell-specific comparisons to the synthetic group. Boxes demarcate the 25th, 50th, and 75th percentile values, while whiskers indicate the outermost point with 1.5 times the interquartile range from the edges of the boxes, white dots indicate mean values. (**b**) Synthetic design methods favor sequences containing heterogeneous program usage. Entropy is calculated from program content to determine homogeneity of program usage for each sequence. For each cell type, synthetic sequences have significantly higher entropy (e.g., lower homogeneity) than all other groups of sequences, except Malinois-natural sequences targeting K562 (Welch's *t*-test p < 1.20e-7). Boxes demarcate the 25th, 50th, and 75th percentile values, while whiskers indicate the outermost point with 1.5 times the interquartile range from the edges of the boxes, white dots indicate mean values. (**c**) Aggregating activating program content, and collapsing over cell types, confirms observations in a. Boxes demarcate the 25th, 50th, and 75th percentile values, while whiskers indicate the outermost point with 1.5 times the interquartile range from the edges of the boxes. (**d**) Same as c, except repressing programs. Boxes demarcate the 25th, 50th, and 75th percentile values, while whiskers indicate the outermost point with 1.5 times the interquartile range from the edges of the boxes, outliers are indicated as points. (**e**) Simultaneous usage of activating and repressing programs and motifs is the favored strategy for synthetic sequence design. Sequences are annotated as activating if composed of at least $2/14^{\text{ths}}$ activating programs and contain one or more motifs with overall positive contributions to activity in the appropriate targeted cell type. Sequences are annotated as repressing if composed of similar repressing program content and contain at least one motif associated with negative contributions in the appropriate off-target cell types. The fraction of sequences in each group passing none, strictly one, or both of these criteria are plotted.

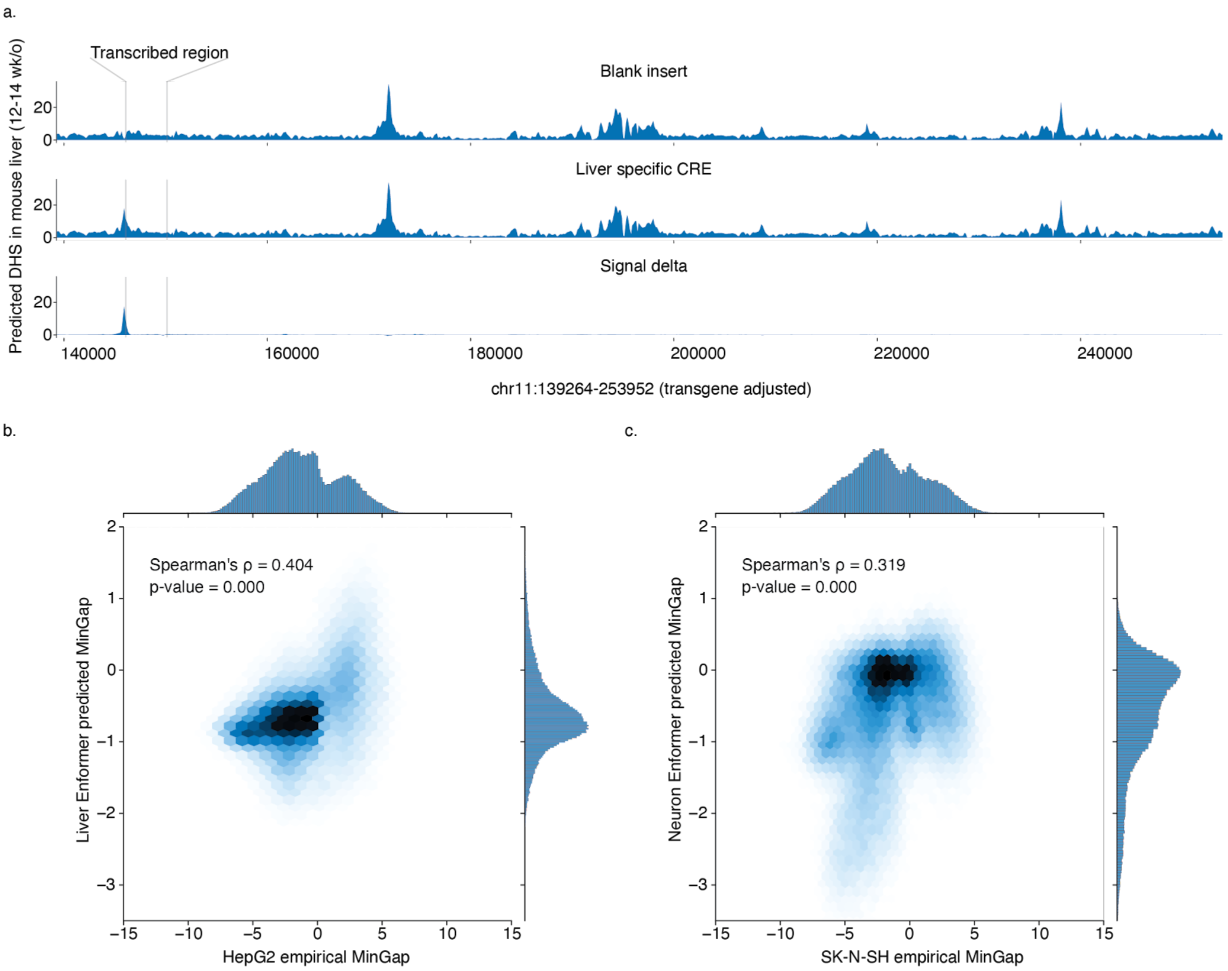


**Supplementary Figure 21. Enformer based prioritization of oligos for *in vivo* tests.** (**a**) Enformer can predict CRE-driven changes in epigenetic and transcription dynamics of transgenes inserted into the H11 safe harbor locus in mice. Three example sequence tracks display predicted DHS signals observed in the livers of 15.5 day old mice. Transgene transcription start site and poly-adenylation signal are indicated by the gray bars. The first track is the predicted signal when the input sequence at the CRE insertion site is all Ns. The second track is an example predicting using a validated HepG2-specific synthetic CRE. The third displays the differential DHS effect. (**b**) Empirical HepG2 MinGap measurements are well correlated with Enformer-predicted features of liver-specific transcriptional activation. (**c**) Empirical SK-N-SH MinGap measurements are also well correlated with Enfomer-predicted features of neural-specific transcriptional activation..

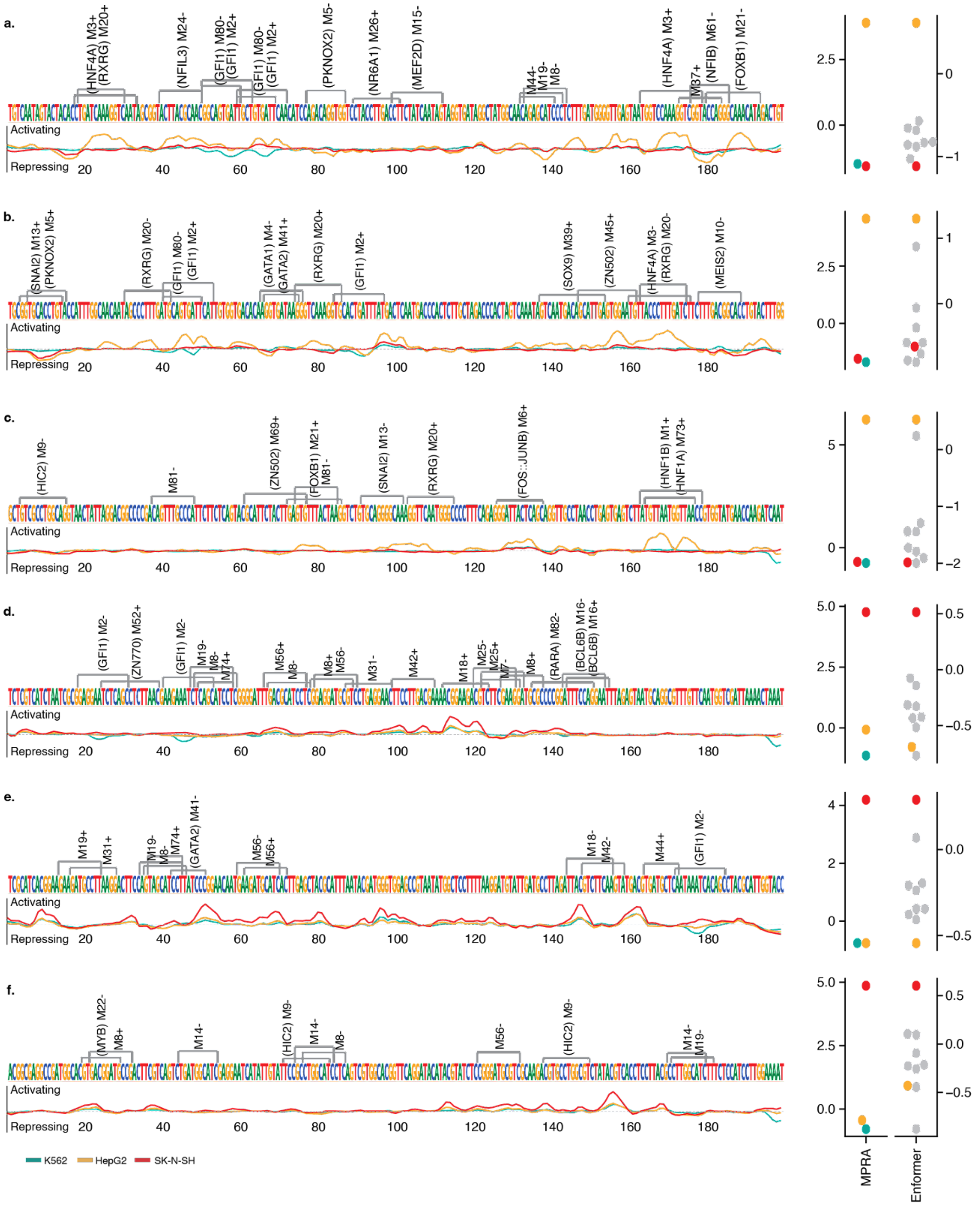


**Supplementary Figure 22. Malinois contribution scores/Enformer/MPRA results for *in vivo* sequences.** Collection of synthetic sequences prioritized for *in vivo* validation. Sequences in panels (**a**-**c**) and (**d**-**f**) are expected to drive expression in liver and neurons, respectively. Left column: Nucleotide sequence, motif matches, and contribution score tracks for each candidate. Right column: Swarm plots of empirical MPRA signal (left *y*-axis) in K562 (teal), HepG2 (gold), and SK-N-SH (red) as well as aggregated Enformer predictions (right *y*-axis) of epigenetic signals reflecting transcriptional activation in mouse liver (gold), neural tissue (red), intestine, kidney, limb buds, lung, pancreas, spleen, and stomach.

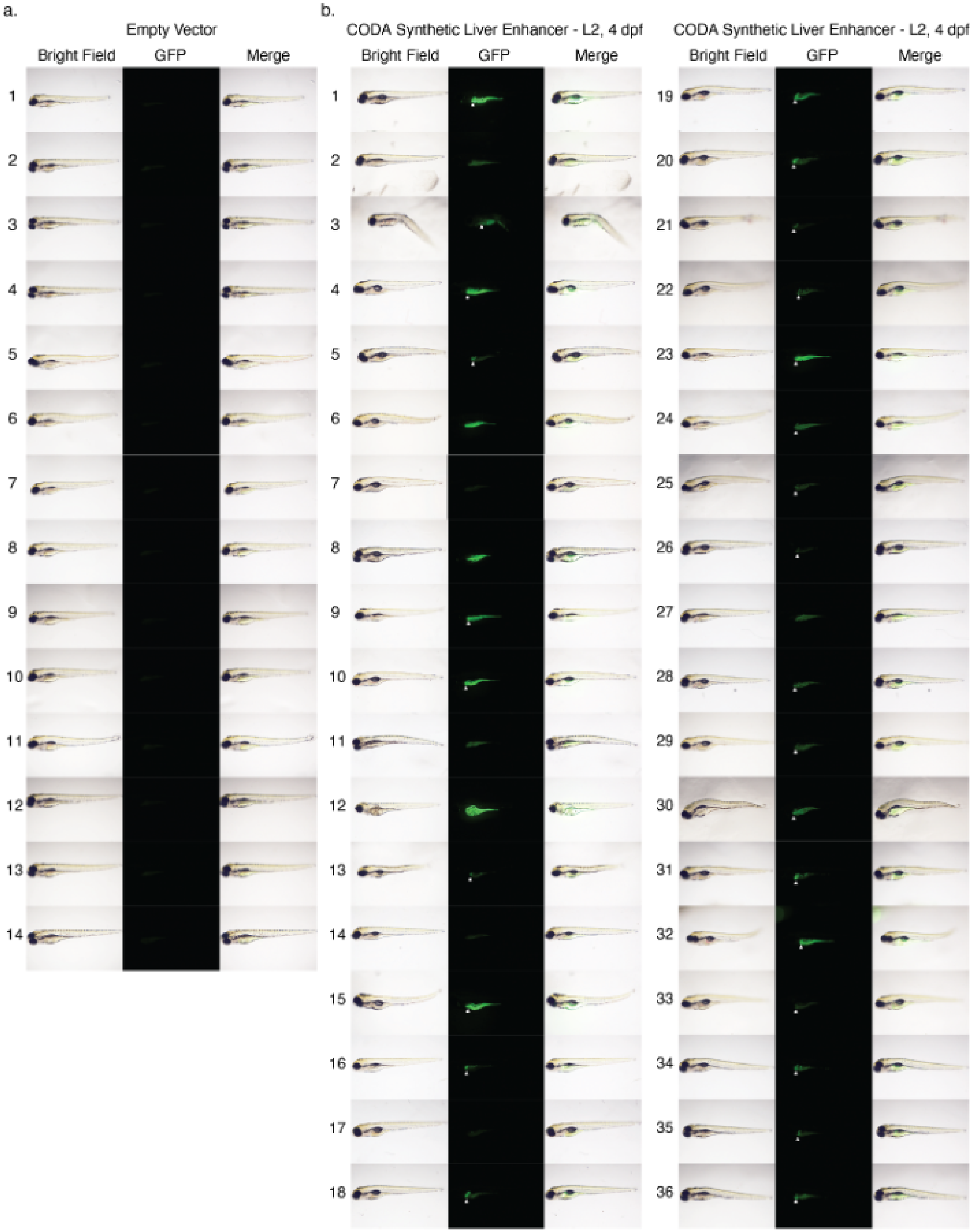


**Supplementary Figure 23. A synthetic CRE reproducibly drives expression in zebrafish livers.** (**a**) Expression of control transgene lacking synthetic CRE fails to drive GFP expression 4 days post-fertilization. All 18 control animals fail to show GFP expression. (**b**) Synthetic CRE drives GFP expression in zebrafish livers and yolk-sacs. Synthetic CRE drives expression in zebrafish livers in 27 out of 36 animals, and yolk-sacs in 32 out of 36 animals.

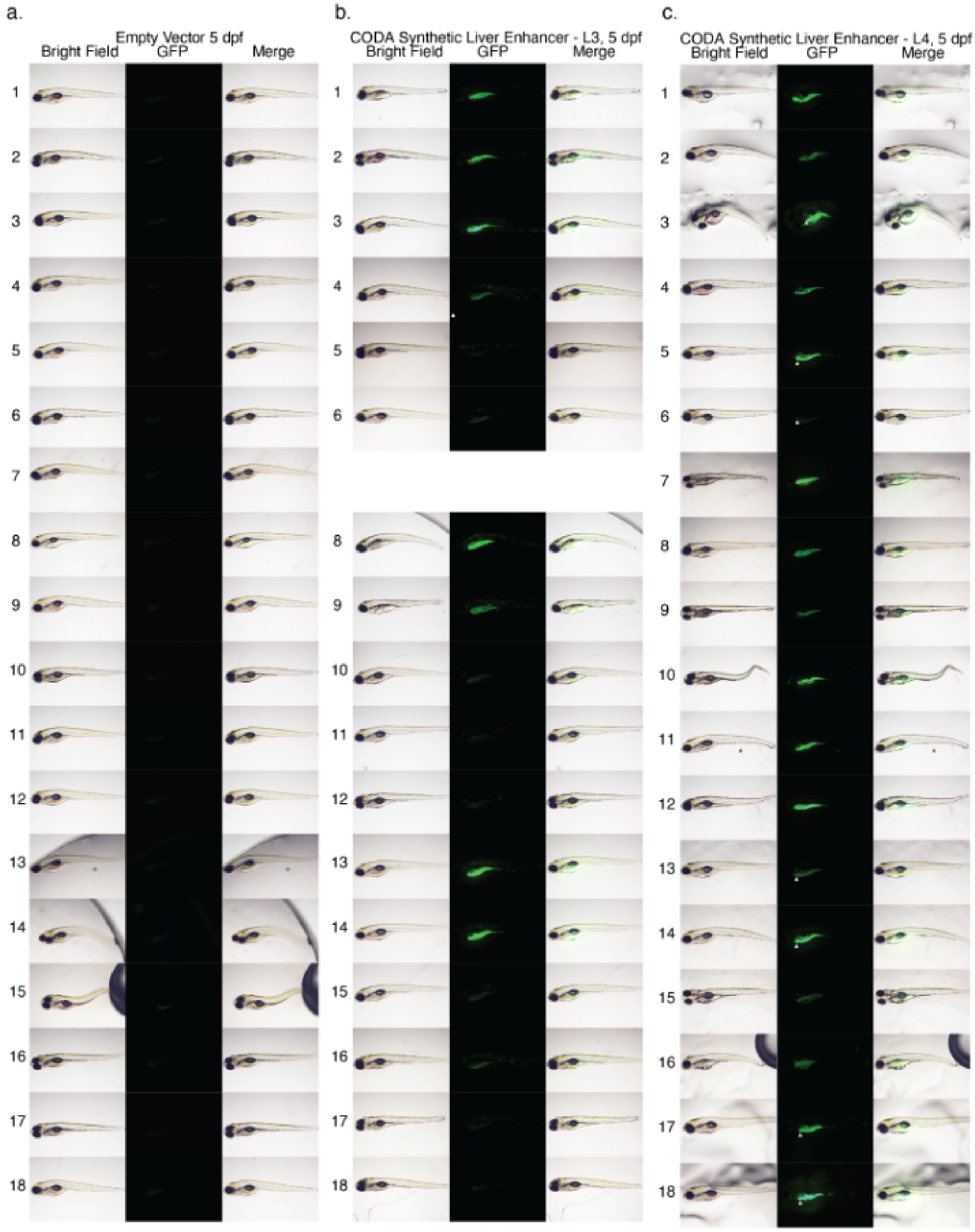


**Supplementary Figure 24. Additional synthetic CREs drive expression in zebrafish gastrointestinal system.** (**a**) Expression of control transgene lacking synthetic CRE fails to drive GFP expression 5 days post-fertilization. All 18 control animals fail to show GFP expression. (**b**) A second synthetic HepG2-specific CRE sporadically drives GFP expression in the yolk-sac, but not the liver. 8 out of 18 animals show CRE induced expression in the yolk-sacs 5 days post fertilization. (**c**) A third synthetic HepG2-specific CRE drives expression drives GFP expression in the yolk-sac.

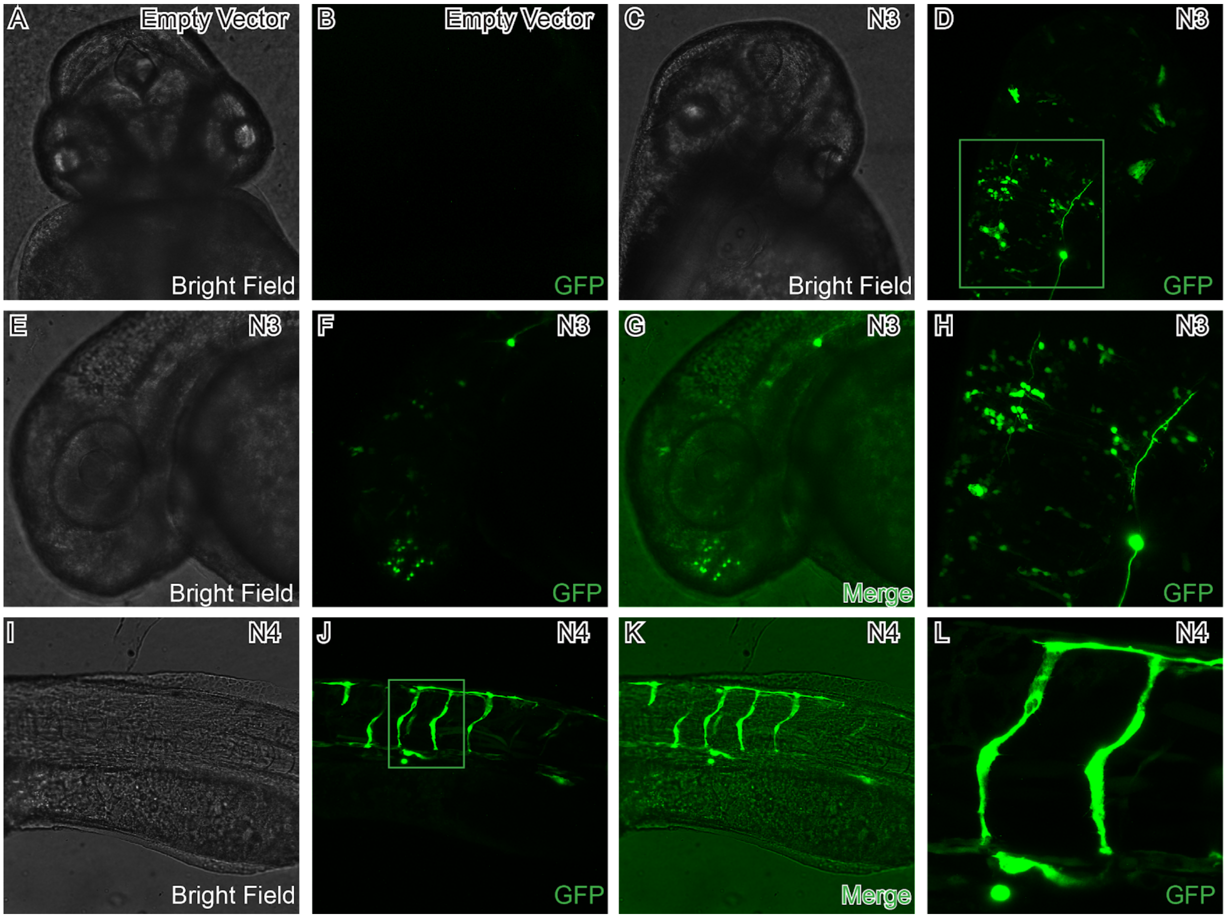


**Supplementary Figure 25. SK-N-SH-specific CREs drive expression in zebrafish neurons or blood vessels.** (**a**) Brightfield image of embryo 48 hours post-fertilization. (**b**) Control transgene lacking synthetic CRE fails to drive GFP expression in head of developing zebrafish. (**c**) Brightfield image of embryo transformed with transgene containing SK-N-SH-specific CRE (N3). (**d**) GFP channel of c. shows transgene expression in neurons. (**e**) Brightfield image of embryo transformed with transgene containing SK-N-SH-specific CRE. (**f**) GFP channel of **e** shows transgene expression in neurons. (**g**) Merged **e** and **f** (**h**) Zoom in of d. (**i**) Brightfield image of embryo transformed with another transgene containing SK-N-SH-specific CRE (N4). (**j**) N4 drives transgene expression in zebrafish blood vessel. (**k**) Merged **i** and **j**. (**l**) Zoom in of **j**. Panels **a**-**d**, **h**: Dorsal views, anterior top. Panels **e**-**g**, **i**-**l**: Anterior to the left, dorsal top.

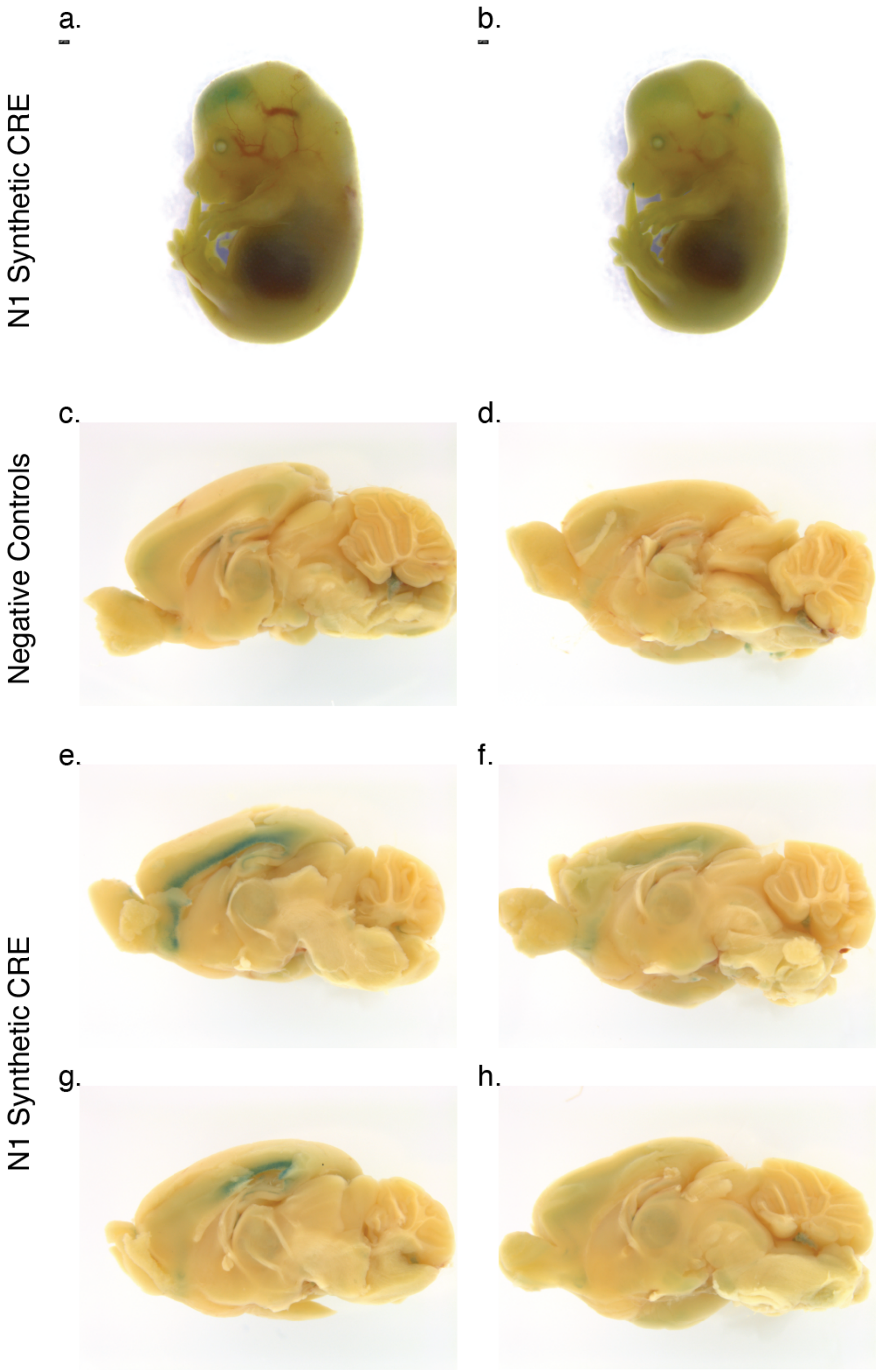


**Supplementary Figure 26. Additional images from mouse transformations.** (**a**) Synthetic neuronal CRE #1 and minP drive transgene expression in developing mouse forebrains. Day 14.5 mouse embryos whole animal lacZ staining. No control mouse. (**b**) Biological replicate of panel **a**. (**c**) Control transgenes with minP-only drive minor transcriptional activation in 5 week old mice. Duplicated from **Figure 4d**. (**d**) Biological replicate of panel **c**. (**e**) Neuronal CRE #1 drives transgene expression cortical layer 6 in 5 week old mouse brains in 3 out of 4 animals. First image is duplicated from **Figure 4d**. (**f**) Biological replicate of panel **e**. (**g**) Biological replicate of panel **e**. (**h**) Biological replicate of panel **e**.

### Supplementary Tables

1 - Ascension information for reference data sets

2 - MPRA Data Used in Training Malinois

3 - Experimental breakdown for CODA library

4 - Penalized motifs in each track/cell type

5 - Motif enrichment (STREME)

6 - Motif matches (FIMO)

7 - Malinois final hyperparameters

8 - Enformer feature indices

9 - Primers and Oligonucleotides

10 - MPRA results and predictions for candidate CRE library
